# Phenotypic characterization of *HAM1*, a novel mating regulator of the fungal pathogen *Cryptococcus neoformans*

**DOI:** 10.1101/2023.09.18.558251

**Authors:** Elizabeth Arsenault Yee, Robbi L. Ross, Felipe H. Santiago-Tirado

**Author notes:** E Arsenault Yee and RL Ross contributed equally to this work, and order was determined by who initiated the study (EAY) and who brought it to completion (RLR). Broad Institute of MIT and Harvard, 415 Main St, Rm 2073, Cambridge, MA 02142.

## Abstract

*Cryptococcus neoformans* is a fungal pathogen responsible for >200,000 yearly cases with a mortality as high as 81%. This burden results, in part, from an incomplete understanding of its pathogenesis and ineffective antifungal treatments; hence, there is a pressing need to understand the biology and host interactions of this yeast to develop improved treatments. Protein palmitoylation is important for cryptococcal virulence, and we previously identified the substrates of its main palmitoyl transferase. One of them was encoded by the uncharacterized gene CNAG_02129. In the filamentous fungus *Neurospora crassa*, a homolog of this gene named HAM-13 plays a role in proper cellular communication and filament fusion. In *Cryptococcus*, cellular communication is essential during mating, therefore we hypothesized that CNAG_02129, which we named *HAM1,* may play a role in mating. We found that *ham1*Δ mutants produce more fusion products during mating, filament more robustly, and exhibit competitive fitness defects under mating and non-mating conditions. Additionally, we found several differences with the major virulence factor, the polysaccharide capsule, that may affect virulence, consistent with prior studies linking virulence to mating. We observed that *ham1*Δ mutants have decreased capsule attachment and transfer but exhibit higher amounts of exopolysaccharide shedding and biofilm production. Lastly, *HAM1* expression is significantly lower in mating media relative to non-mating conditions, consistent with it acting as a negative regulator of mating. Understanding the connection between mating and virulence in *C. neoformans* may open new avenues of investigation into ways to improve the treatment of this disease.

**Importance:** Fungal mating is a vital part of the lifecycle of the pathogenic yeast *Cryptococcus neoformans*. More than just ensuring the propagation of the species, mating allows for sexual reproduction to occur and generates genetic diversity as well as infectious propagules that can invade mammalian hosts. Despite its importance in the biology of this pathogen, we still do not know all of the major players regulating the mating process and if they are involved or impact its pathogenesis. Here we identified a novel negative regulator of mating that also affects certain cellular characteristics known to be important for virulence. This gene, which we call *HAM1*, is widely conserved across the cryptococcal family as well as in many pathogenic fungal species. This study will open new avenues of exploration regarding the function of uncharacterized but conserved genes in a variety of pathogenic fungal species, and specifically in serotype A of *C. neoformans*.

## 1. Introduction

There are approximately 4 million fungal species worldwide (1), and although most play beneficial roles in their environments, a few hundred are known to be pathogenic to mammalian hosts (2). A common theme in these fungal infections is the importance of morphological transitions for pathogenesis. *Histoplasma capsulatum* grows naturally as a mold in the environment but under host conditions, it grows as a budding yeast which allows for greater dissemination in the bloodstream (3). Conversely, *Aspergillus fumigatus* conidia are inhaled into the lungs and germinate into hyphal filaments, which can form mycelial networks that are able to invade into host tissues and due to their large size, are unable to be phagocytosed by host immune cells. In the case of *Cryptococcus neoformans*, one such morphological transition happens prior to host invasion, where sexual and asexual reproduction produces the spores and desiccated yeasts that are the infectious agents inhaled into the lungs to establish infection (4).

*C. neoformans* is an opportunistic basidiomycete pathogenic fungus present worldwide (4). Because it is in the environment, the infectious propagules are frequently inhaled into the lungs, causing an infection that in healthy individuals is controlled and remains asymptomatic until cleared. However, in the immunocompromised, the infection quickly overwhelms the immune system in the lungs, resulting in pneumonia and acute respiratory distress syndrome, with subsequent dissemination to the central nervous system, causing lethal meningoencephalitis (5). *C. neoformans* is a leading cause of death in HIV/AIDS, where there are approximately 152,000 cases each year with 112,000 resulting in death (6). This staggering mortality rate stems largely from a lack of effective treatment options as well as an incomplete understanding of cryptococcal pathogenesis. This has prompted the World Health Organization to include *C. neoformans* in the highest critical priority group of its recent Fungal Priority Pathogen List (7).

In a previous study investigating cryptococcal factors necessary for phagocyte recognition and internalization (8), we identified a gene homologous to *Saccharomyces cerevisiae*’s Protein Fatty Acyltransferase 4 (*PFA4*). We and others showed that cryptococcal *PFA4* was necessary for complete virulence in a mammalian host (8, 9). Since palmitoyl acyl transferases (PATs) like Pfa4 act by modifying substrate proteins, we determined its palmitoylome. Using comparative mass spectrometry between wild-type (WT) and *pf*a*4*Δ samples to determine downstream targets, one of the most highly enriched substrates was encoded by the gene of unknown function CNAG_02129 (8).

CNAG_02129 is part of a conserved family of fungal genes of which only one member, from the filamentous ascomycete *Neurospora crassa*, has been partially characterized (hyphal anastomosis protein 13 or HAM-13)(10)(Fig. 1A). In the absence of HAM-13, only 33% of the *N. crassa* germlings find each other and fuse. This is in stark contrast to almost 100% of the WT germlings finding each other, undergoing cellular fusion, and maturing into interconnected hyphae to form a mycelium (11). Thus, the authors concluded that HAM-13 plays a role in cellular communication. In *C. neoformans*, this type of cellular communication is essential for mating, which can be either unisexual or bisexual depending on the surrounding environmental pressures such as nutrient limitation or pheromone secretion (12). Members of the *C. neoformans* species complex (which includes seven pathogenic species) have two mating types determined by the *MAT* loci, *MAT*α and *MAT*a. In a bisexual scenario, *MAT*α and *MAT*a cells find each other through pheromone signaling, which triggers a cell fusion event resulting in a morphological transition into filamentous hyphae. The nucleus from each parent travels up these hyphal structures to the apex where a basidia is formed. Within the basidia, nuclear fusion occurs followed by meiosis and several rounds of mitosis to produce four chains of haploid basidiospores. These spores then drop from the basidium and mature into yeast where the process can be started again. In a unisexual scenario, the same events occur but only in cells containing the *MAT*α locus (12). Almost all of these details, however, are from studies using serotype D *C. neoformans* (now called *C. deneoformans*), and considerably less is known about *C. neoformans* strains that are serotype A (*C. neoformans sensu stricto*).

**Figure 1:**
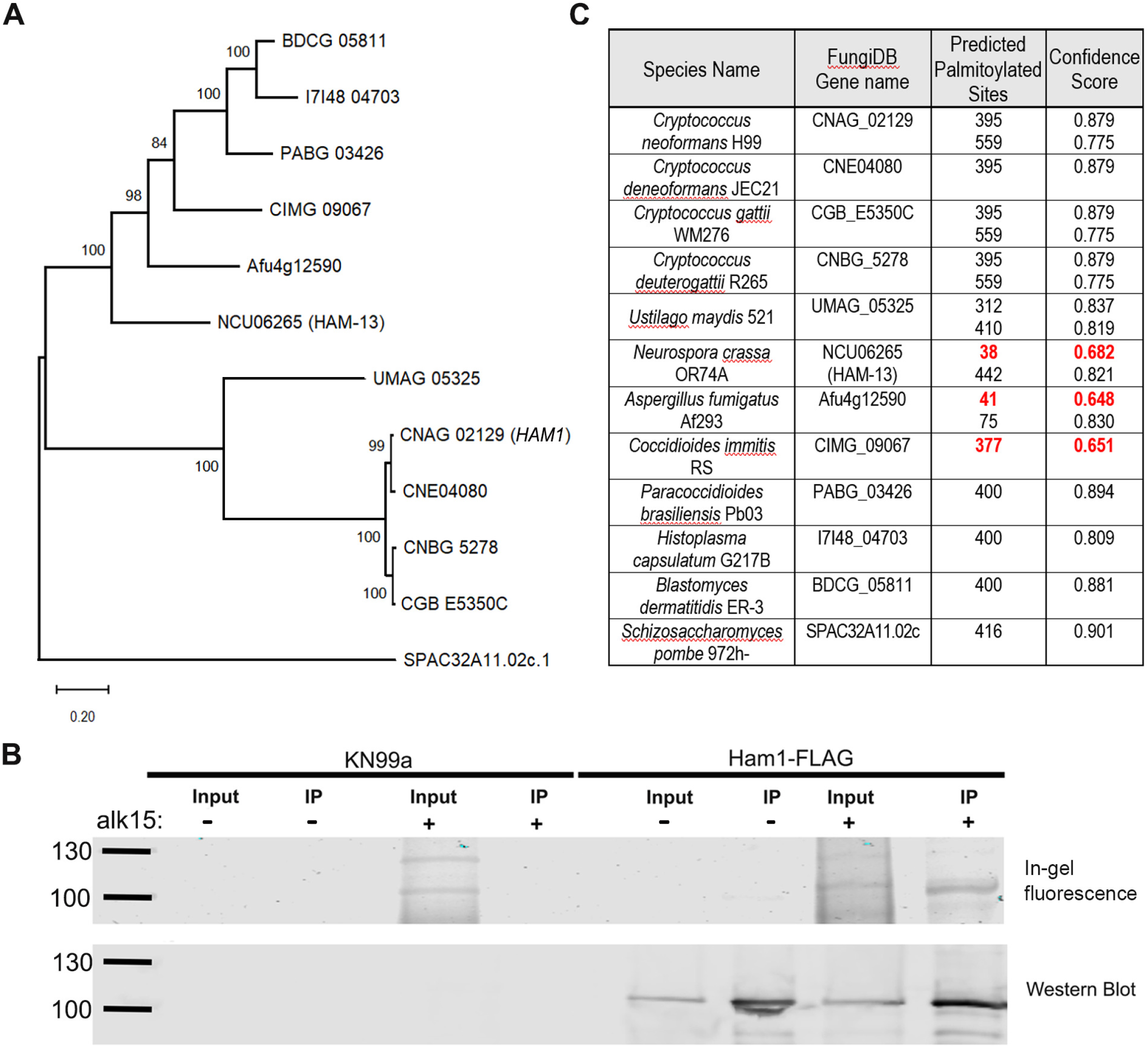
Ham1 is a palmitoylated protein conserved across the fungal kingdom. (A) Phylogenetic tree showing conserved homologs of *HAM1* (CNAG_02129) across common pathogenic and model fungal species. The evolutionary history of the sequences was inferred using the Maximum Likelihood method with MEGAX software (48). The tree with the highest log likelihood (-11862.20) is shown. 1,000 replicate analyses (bootstraps) were run and the percentage of trees in which the associated taxa clustered together is shown next to the branches. The tree is drawn to scale, with branch lengths measured in the number of substitutions per site. (B) Immunoprecipitation (IP) coupled with bioorthogonal click chemistry to probe palmitoylation of Ham1. Cultures of untagged (KN99) or tagged (Ham1-FLAG) cells were either not fed (–) or fed (+) an analog of palmitate (alk15) and then lysed to generate a protein extraction. An IP was then performed on fed and unfed fractions to pull down Ham1-FLAG which was then coupled to a fluorescent reporter by click chemistry, highlighting palmitoylated proteins. (Top row) A representative in-gel fluorescence experiment showing signal only in the fed fractions. (Bottom row) The accompanying western blot of the same samples run above, probed with anti-FLAG antibody. Ham1-FLAG is ∼102 KDa, highlighted ladder lanes are 100 and 130 KDa. (C) Predicted palmitoylation scores of *HAM1* and homologous genes used in (A), using GPS-Palm (17). Residues predicted with low confidence (scores 0.6484 – 0.7750) are highlighted in red.

Several studies have shown an inverse relationship between *C. neoformans’s* virulence and its filamentous mating form (13, 14). For example, when *ZNF2,* a master regulator of filamentation in *C. neoformans*, is constitutively overexpressed, the cells are morphologically hyphal-locked and unable to cause disease in a mouse model (15). These hyphal-locked cells induced a protective TH-1 pro-inflammatory immune response that not only resulted in full clearance of the initial infection, but also protection from subsequent challenges (16). This suggests that mating regulators may also play a larger role in control of virulence mechanisms, such as impacting any of the main cryptococcal virulence factors: its polysaccharide capsule, thermotolerance, and melanization. These and other factors allow for *C. neoformans* to persist within the harsh environment of a human host. While each of these virulence factors has been comprehensively studied, not all their regulators have been uncovered.

In this study, we determined that CNAG_02129, which we refer to as *HAM1* (Hyphal anastomosis protein 1), plays a role in cryptococcal mating by acting as a negative regulator of the process, and has a connection to proper polysaccharide capsule retention and shedding. Our *ham1*Δ strains exhibit increased cellular fusion with and without exogenous pheromone, a hyper hyphal response during bisexual mating scenarios, and reduced competitive fitness under both mating and non-mating conditions. Interestingly, these effects were specific for *C. neoformans* (serotype A) and were not obvious in *C. deneoformans* (serotype D), highlighting the known differences in mating regulation between the sister species. Notably, although *HAM1* expression is significantly downregulated during mating conditions, there are no differences in *MAT* loci pheromone expression over time, suggesting a fork somewhere in the pheromone signaling cascade. Consistently, *HAM1* expression still is high under mating conditions in sterile mutants (*cpk1*Δ and *mat2*Δ) that have no pheromone expression. Although we do not observe major defects in capsule generation and other key virulence mechanisms, we found defects in capsule attachment, an increase in capsule shedding, less effective capsule transfer, and an increase in biofilm production. These findings suggest a connection between mating and proper capsule formation and shedding, which can impact the virulence potential of this organism, and contribute to our understanding of mating regulation in this important fungal pathogen.

## 2. Results

### 2.1 Ham1 is a palmitoylated protein conserved across the fungal kingdom

To confirm the results of our previous large scale Pfa4 palmitoylome studies, we investigated the palmitoylation status of Ham1 using a click-chemistry-based palmitoylation probe (8). Using this probe (alk15; an analog of palmitate) and a *HAM1*-FLAG tagged strain (see Methods), we performed an immunoprecipitation (IP) coupled with click-chemistry to determine if the Ham1 protein was palmitoylated. We confirmed that Ham1 is palmitoylated by the presence of a fluorescent band only in the probe-fed fraction of our Ham1-FLAG IP that was at the same size as the accompanying western blot (Fig. 1B). Next, we wanted to see if the palmitoylation modification may be conserved across the protein homologs. Using the palmitoylation predictor software GPS-Palm (17), we determined that all of the homologs investigated had very high confidence predictions for palmitoylation (Fig. 1C).

### 2.2 *ham1*Δ exhibits altered cellular fusion and progeny with a dry colony morphology

The goal of mating is to produce diploids that can undergo meiosis and generate haploid cells with traits from both parents, generating better-adapted cells (12). This requires that parental strains find each other, fuse, undergo recombination, and then produce haploid spores. To test if mating was affected, deletion strains of *HAM1* were generated in both mating types of the KN99 background (WT) using different dominant markers (NEO or NAT). We then set up unilateral (one mating partner is mutant, the other is WT) and bilateral (both mating partners are mutant) mating crosses between WT mating controls (KN99-NEO or KN99-NAT) and *ham1*Δ mutants (*ham1*Δ-NAT), and determine the fusion efficiency from the number of colonies resistant to both antibiotics. Both unilateral and bilateral *ham1*Δ mutant crosses yielded significantly higher colonies resistant to both antibiotics when compared to the WT cross, indicating faster, or more efficient, cellular fusion (Fig. 2A, Supplemental Fig. 1). Notably, the *ham1*Δ mutants are still under pheromone regulatory control, as there are no fusion events detected between the same mating types (Supplemental Fig. 1A). However, to explore the impact that pheromone might have on cellular fusion, we tested if adding synthetic exogenous pheromone would increase the rate of cell fusion in our *ham1*Δ mutants (Fig. 2A). Although addition of exogenous pheromone to the WT cross resulted in an increase in the number of colonies generated, there was no significant difference in the fusion rate (Supplemental Fig. 1B). Likewise, in our *ham1*Δ crosses there was no statistical significance by one-way ANOVA with multiple comparisons (Supplemental Fig. 1C, D), indicating that this enhanced fusion is pheromone-signal independent. Interestingly, when scoring double-resistant colonies, we observed a difference in their morphology. Rather than the typical glossy, round shape colonies coming from the WT strain crosses, the fusion products from the *ham1*Δ crosses had a dry, irregular morphology (Fig. 2B). When resuspending single colonies from these crosses and staining them with DAPI and CFW to visualize, we found that these colonies were comprised of the hyphal form of *Cryptococcus,* as the septal divisions and a binuclear distribution along septa were evident (Fig. 2B). In contrast, the colonies from the WT crosses only showed budding yeasts. This drastic increase in the ability to fuse as well as the dry appearance and hyphal state of *ham1*Δ progeny led us to consider what other facets of the mating cycle could be altered.

**Figure 2:**
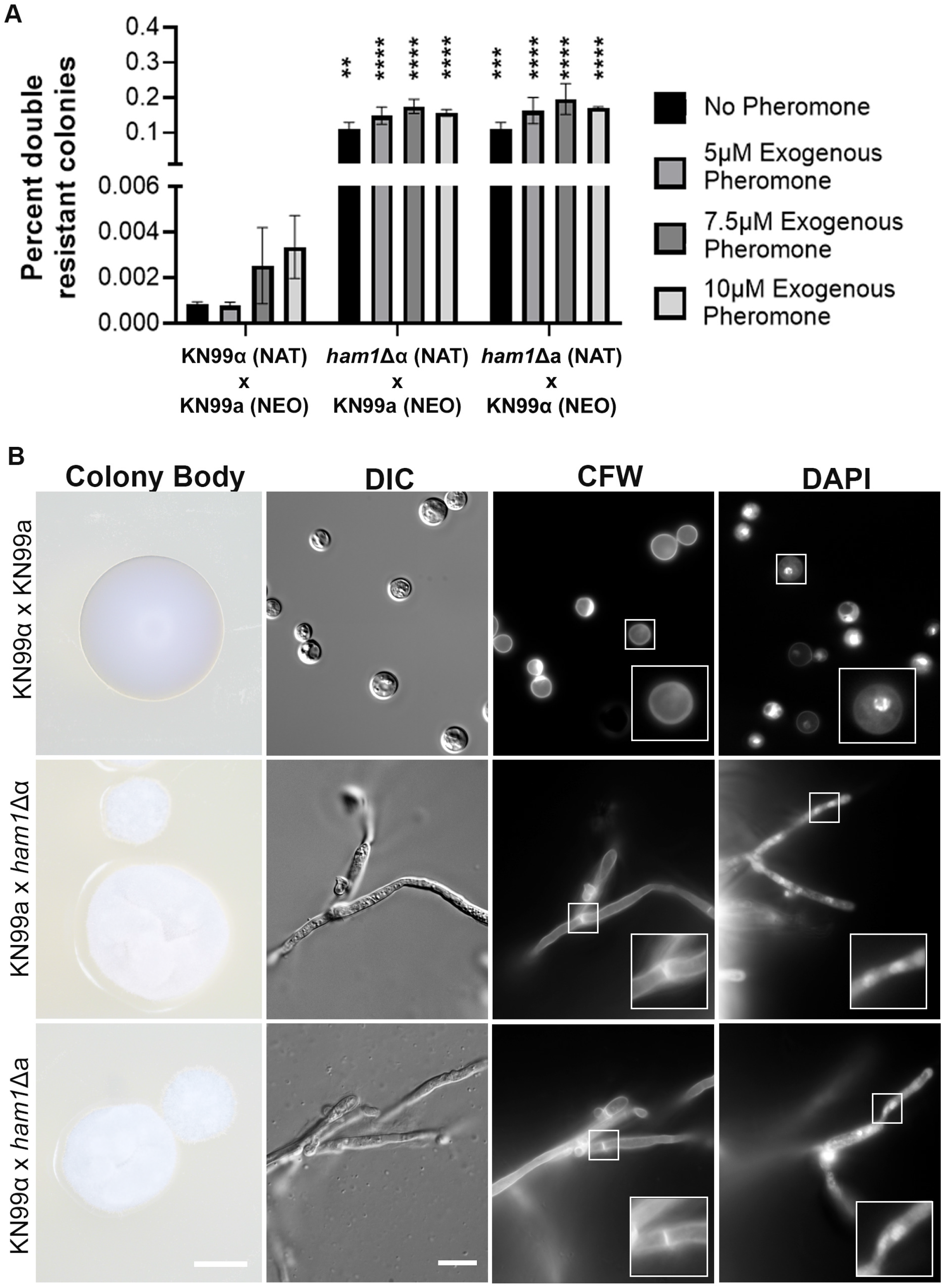
*ham1*Δ mutants have higher cellular fusion efficiency and exhibit a dry colony morphology. (A) The percent of colonies growing in double antibiotic plates (# of colonies that are NAT and G418 resistant over total colonies) resulting from the indicated bisexual crosses is shown. Each condition (no pheromone and with the different concentrations of exogenous pheromone) was compared to the WT controls (KN99α x KN99a) by 2-way ANOVA with a Dunnet’s multiple comparison test; ** P < 0.01, *** P < 0.001, **** P < 0.0001; N = 3 biological replicates for all crosses and all conditions. (B) Representative images of colony morphology (Colony Body) from all crosses at 2x magnification, and 100x magnification of cells from each colony stained with Calcofluor white (CFW) to highlight septal divisions of hyphal form in mutant crosses, and with DAPI to show binuclear distribution in mutant crosses. Scale bars represent 500 and 10 microns, respectively.

### 2.3 *ham1*Δ mutants have increased hyphal production in bisexual mating scenarios on V8 medium

Hyphal growth is a quantitative trait that can be used to measure how well a strain is able to mate (18). We assessed the ability of our *ham1*Δ crosses to form hyphae in a time course from 5 days to 4 weeks. We observed faster and more robust filamentation in our *ham1Δ* crosses across all time points (Fig. 3A). When quantifying the hyphal area (see Methods), we observed a significant increase in all unilateral mutant crosses at 5 days and 3 weeks, and a significant increase in the hyphal area of the bilateral crosses across all time points (Fig. 3B). This was confirmed using two additional independent *ham1Δ* mutants, including one from the deletion collection available commercially (www.fgsc.net; Supplemental Fig. 2A).

**Figure 3:**
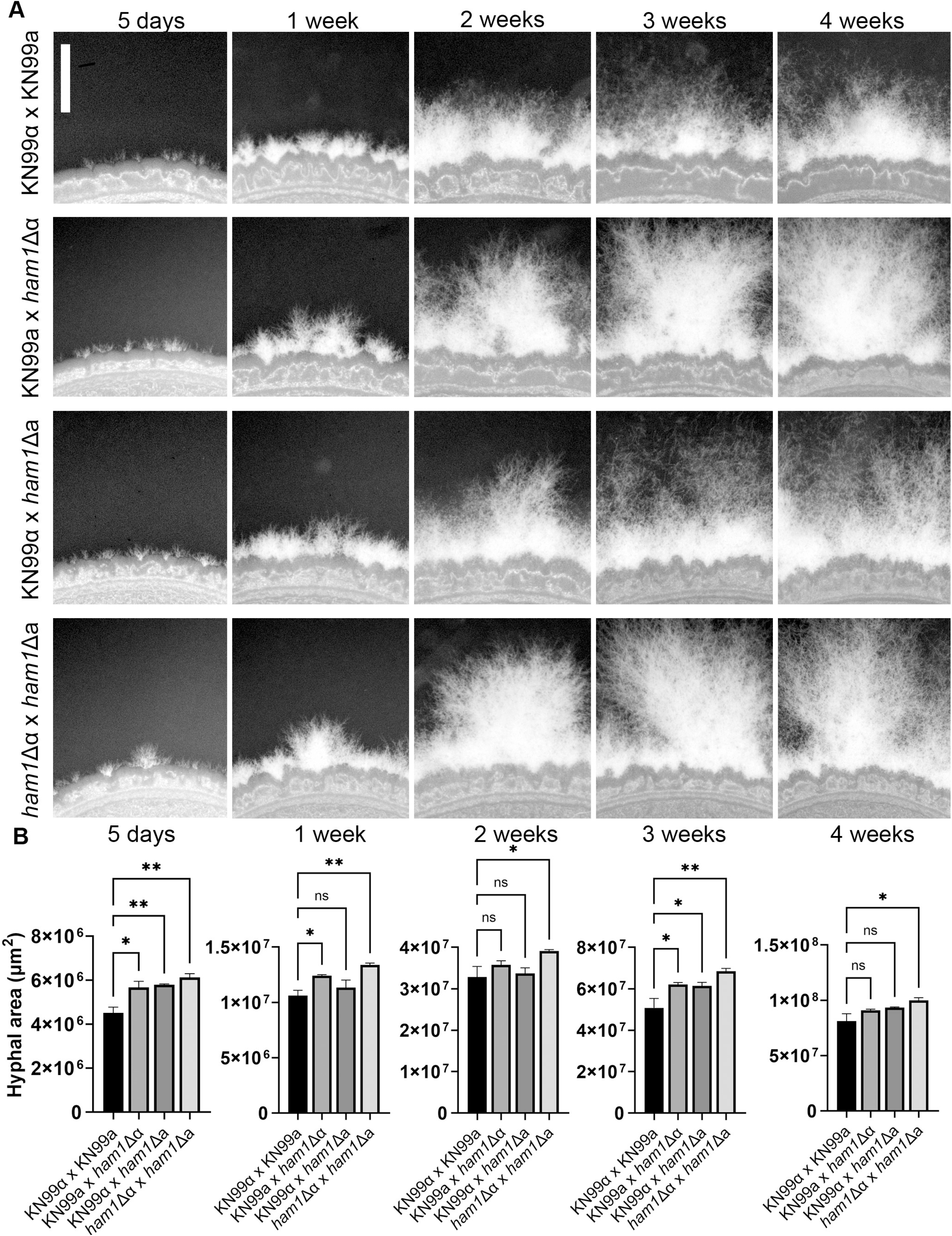
*ham1*Δ unilateral and bilateral crosses exhibit increased hyphal production on V8 mating medium. (A) Representative images of WT and mutant crosses on V8 mating media. All images taken at 63x magnification scale; bar represents 1 mm. (B) Quantification of hyphal area for all crosses at all time points. Hyphal area was calculated in FIJI (ImageJ) as the difference between a cell body mask and a mask covering the whole cell body and surrounding filaments. All mutant crosses were compared to WT controls (KN99α x KN99a) by one-way ANOVA with Dunnet’s multiple comparison test; * P < 0.05, ** P < 0.01. N = 3 biological replicates for all timepoints of hyphal area.

### 2.4 *ham1*Δα strain exhibits a fitness disadvantage in both mating and non-mating scenarios

Mating is an energy costly process, and may result in a fitness cost when comparing mating and non-mating conditions. However, this is crucial for survival in the environment where mating partners are few and far between and the pressures of predators and other stressors threaten survival. We tested if our *ham1*Δ would display increased fitness compared to a WT competitor due to its propensity to form more hyphae and increased cellular fusion events. To test this, we had two *MAT*α or two *MAT*a strains with opposing resistance cassettes (the competitors) co-incubated with the opposing mating type with no resistance cassette (the donor) (Fig. 4A). All three strains were mixed and plated under mating (V8) or non-mating (YPD) conditions for 10 days. At that point, each competitive scenario was scraped, resuspended, and plated on selective media to determine the colony forming units (CFUs) of each competitor and calculate a competitive index (CI; Fig. 4B). In the mating permissive condition (V8) we expect both competitors to stop vegetative budding and forage for mating partners while in the non-mating condition (YPD) we expect both competitors to simply divide. In the WT cross we observed no fitness defects (CI ≈ 1) in either YPD or V8 media conditions (Fig. 4C). However, in our *ham1*Δ mutant crosses we saw CI values significantly different from 1. In the *ham1*Δα competition, the CI was less than one (0.17 in YPD and 0.32 in V8), indicating significant fitness defects for *ham1*Δα in both the V8 and YPD conditions (Fig. 4C). Surprisingly, in our *ham1*Δa competition we observed the opposite, CI values larger than one (1.84 in YPD and 1.65 in V8) showing a strong competitive advantage in both conditions for *ham1*Δa (Fig. 4C).

**Figure 4:**
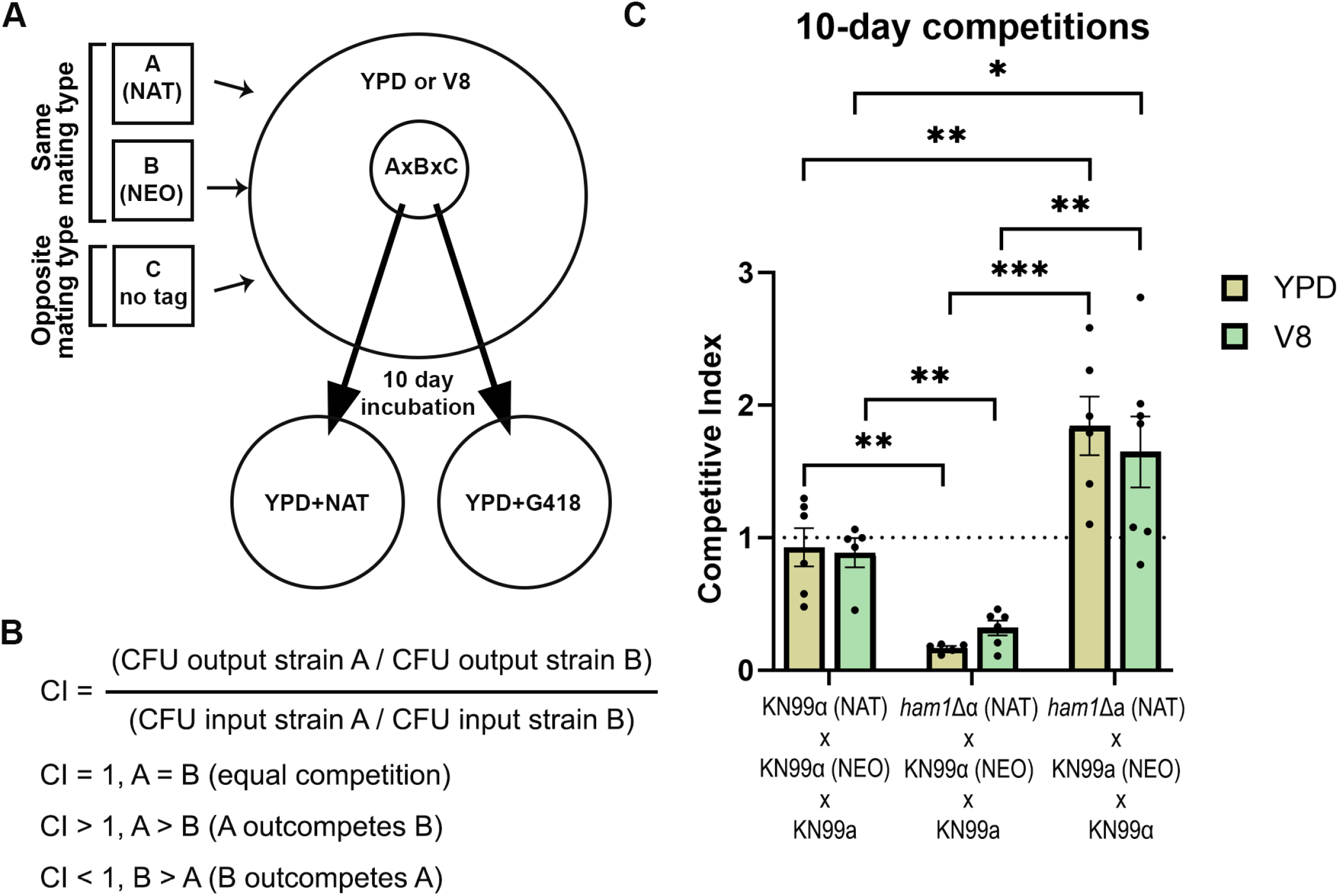
*ham1*Δ mutants have altered competitive fitness when compared to WT. (A) schematic of how the cellular competition assay was performed. (B) Definition of the competitive index (CI) used to compare the fitness of strain A and strain B in the mixture of three strains. (C) Outcomes of the competitive assays for each of the indicated mixtures. The dashed horizontal line represents a CI of 1, or equal competition. Black circles represent the average of each independent assay, while the bar shows the mean and standard error of all assays (n = 5 - 7 biological replicates for all competition assays). Values were compared by one-way ANOVA with uncorrected multiple comparisons (unpaired t tests with Welch’s correction). * P < 0.05; ** P < 0.01; *** P < 0.001.

### 2.5 Transcriptional analysis of the pheromone pathway and *HAM1* over time

To determine if *HAM1* is involved in the pheromone response MAP kinase pathway we analyzed the transcriptional activity of the *MAT*α/*MAT*a pheromone loci and *HAM1* under YPD and V8. This type of analysis is typically done in *C. deneoformans*, but when we deleted *HAM1* in this species, mating was unaffected (data not shown). Since it is known that there are differences in mating regulation between the two sister species, we decided to continue and analyze the transcription of these genes in *C. neoformans*. To do this for the pheromone loci, we compared the gene expression of *MAT*α and *MAT*a in mating conditions normalized to the expression in non-mating conditions to get the relative fold induction (19, 20). We performed this in a time course of 1, 3, 5 and 7 days to look at changes in expression over time (Fig. 5 A-C). Not surprisingly, we observed no statistically significant differences in either *MAT*α or *MAT*a expression between the crosses. However, in the WT cross, we saw a strong and consistent downregulation of *HAM1* under mating conditions relative to non-mating media. We saw that on day 1 in mating conditions, *HAM1* had a much lower expression as compared to non-mating conditions (Fig. 5C). However, as the mating cycle progressed, the expression of *HAM1* returned to levels similar to those under non-mating conditions (Fig. 5C). These findings, taken together with the increased fusion rate and filamentation of the *ham1*Δ mutants, suggest that *HAM1* acts as a negative regulator of mating under non-mating conditions. This would imply that the pheromone MAP kinase pathway downregulates expression of *HAM1* to promote mating. To test this, we measured *HAM1* expression in WT, *cpk1*Δ, and *mat2*Δ mutants under both mating and non-mating conditions (Fig. 5D). Consistently, in bisexual crosses between either of these mutants (known to be sterile mutants; (21)) and KN99a, *HAM1* expression remains high. The same happens when only one mating partner is grown in YPD or V8 (both would be non-mating conditions for *C. neoformans,* as this species does not undergo unisexual reproduction), the relative *HAM1* expression is even higher in V8 than in YPD. Interestingly, when measuring *HAM1* expression in monocultures of *cpk1*Δ or *mat2*Δ, the *mat2*Δ cells still greatly induce *HAM1* in V8 relative to YPD just like the WT monocultures, whereas in the *cpk1*Δ monoculture *HAM1* was downregulated similar to WT bisexual crosses.

**Figure 5:**
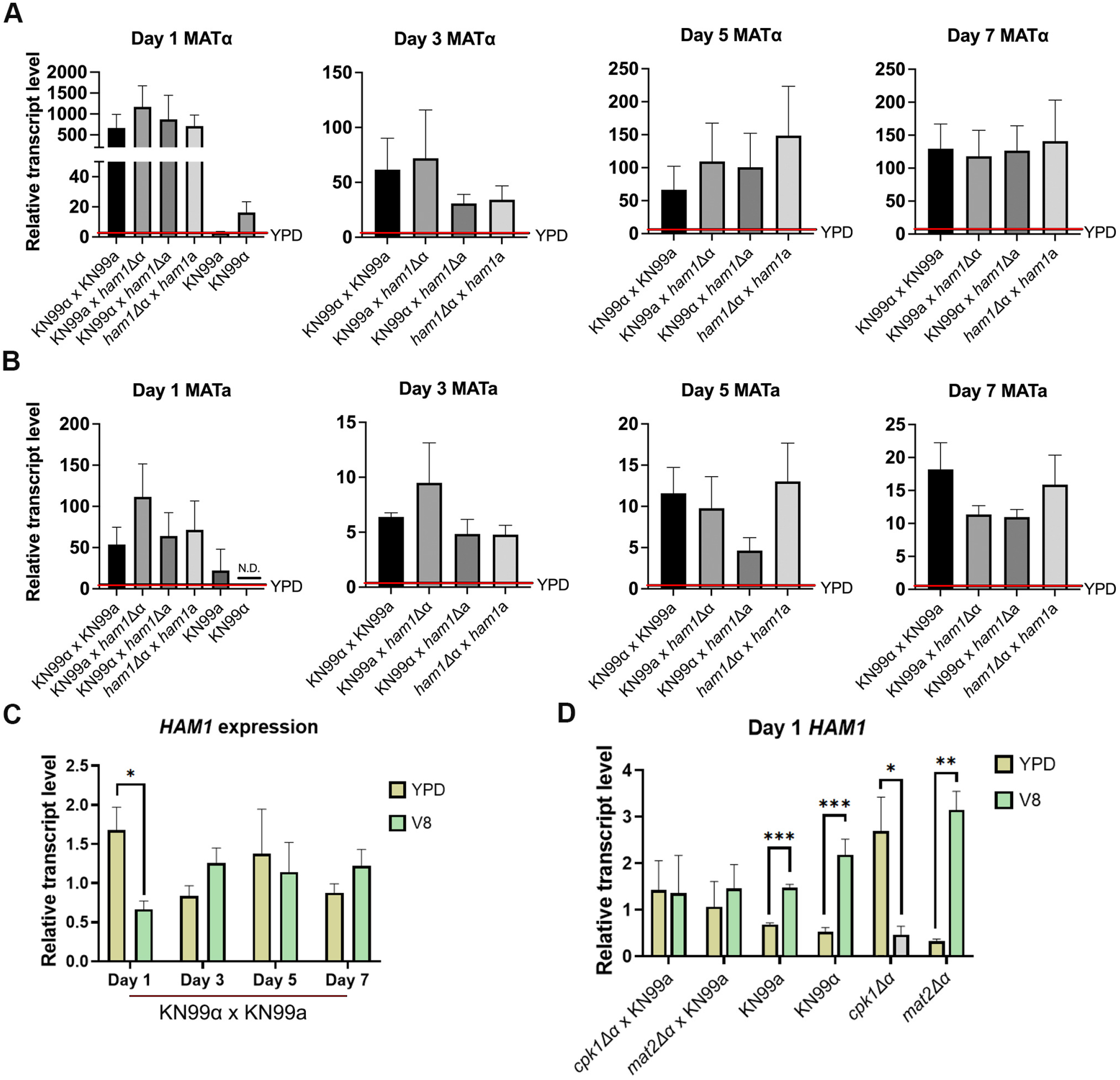
*HAM1* does not regulate pheromone expression but its expression correlates with the mating status of the cell. (A) Relative transcriptional analysis of *MAT*α in mating conditions (V8 media) relative to non-mating conditions (YPD media) after incubation for 1, 3, 5, and 7 days on the respective media. (B) Relative transcriptional analysis of *MAT*a in mating conditions (V8 media) relative to non-mating conditions (YPD media) after incubation for 1, 3, 5 and 7 days on the respective media. N.D., not detected. (C) Quantification of the transcript level of *HAM1* in non-mating (YPD) and mating (V8) conditions after incubation for 1, 3, 5 and 7 days on each media. Each day, the YPD values were compared to the V8 values by a Mann-Whitney test. * P < 0.05. N = 5 biological replicates for all timepoints and conditions. (D) Same as (C), but measured in the indicated crosses or monoculture suspensions. For each cross, or monoculture suspension, the YPD values were compared to the V8 values by a Mann-Whitney test. * P < 0.05; ** P < 0.01; *** P < 0.001. N = 2 biological replicates for all timepoints and conditions. Red line shows expression levels in YPD.

### 2.6 *ham1Δ* has no significant defects in major virulence mechanisms

So far, all the phenotypes of the *ham1*Δ have been mating-related, especially in the *MAT*α background. We wondered if these would also affect virulence; hence, we investigated if there were defects in the main virulence factors of *C. neoformans*: capsule production, melanization, thermotolerance (37°C), and recognition by phagocytic cells. We found no obvious differences in any of the major virulence factors (Fig. 6, Supplemental Fig. 3, and data not shown). Due to the surface defects observed in the *pfa4*Δ mutant (Ham1’s PAT)(8) we took a closer look at the cell wall and the capsule of our *ham1*Δ strains as more nuanced differences could be present besides capsule induction. Finding no apparent defects under different cell wall and membrane stressors (Supplemental Fig. 3), we took a closer look at the polysaccharide capsule and investigated differences in its attachment and release.

**Figure 6:**
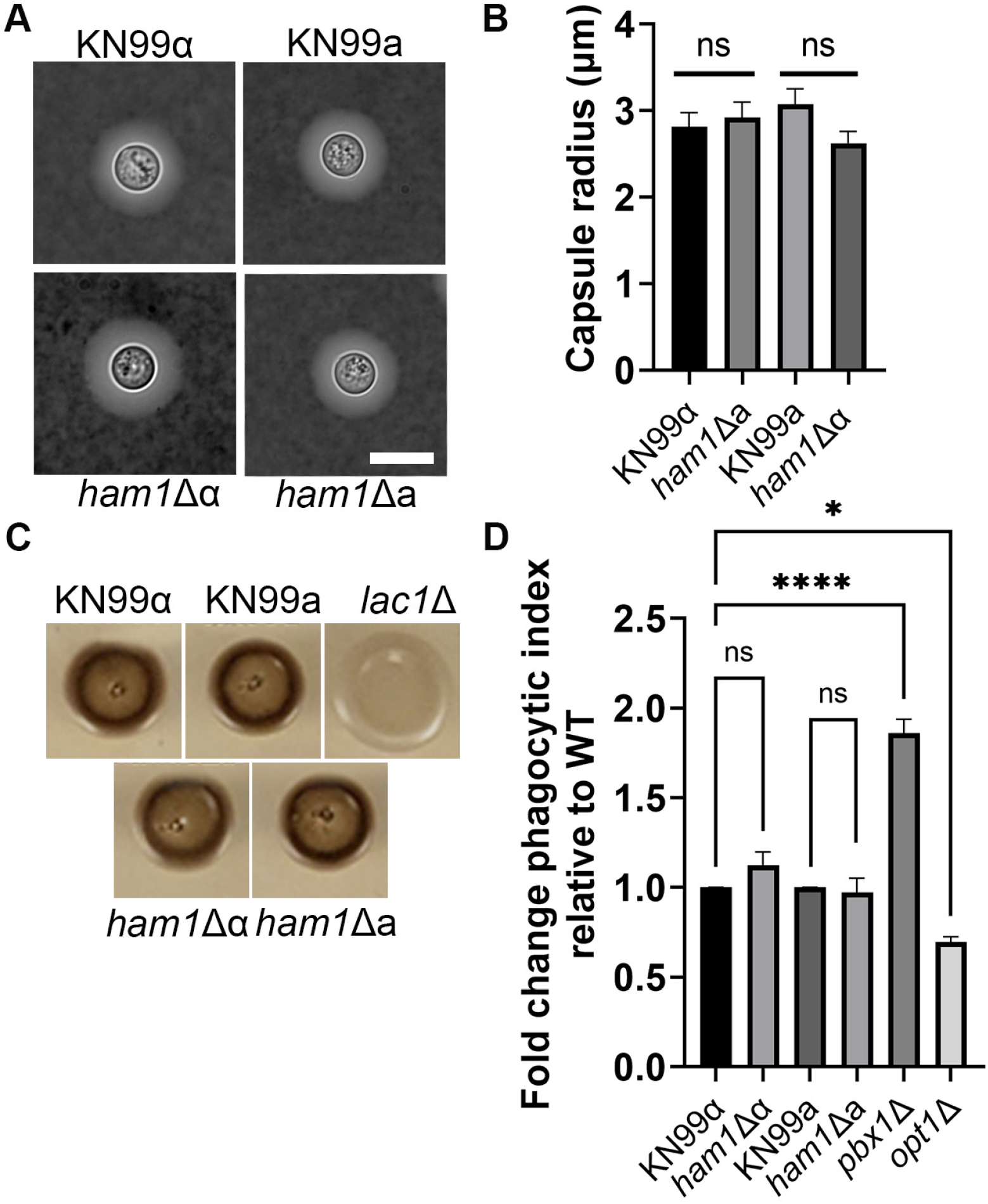
*ham1*Δ shows no defects in major virulence mechanisms. (A) Representative images of India Ink staining of induced capsule in the indicated strains. Scale bar represents 10μm. (B) Quantification of capsule size after induction. Shown is the average value; N = 25 cells for all strains. (C) Indicated strains were spotted on caffeic acid plates to visualize melanin production; *lac1*Δ is a melanin mutant and serves as a negative control. (C) Measurement of uptake by THP-1 phagocytic cells as a function of the number of engulfed fungi divided by the number of THP-1 cells, normalized to the WT controls; *pbx1*Δ is a known mutant with a higher uptake index and *opt1*Δ is a known low uptake mutant. Results were compared by a one-way ANOVA with Dunnet’s multiple comparison test for all strains with the *MAT*α background, this includes our positive (*pbx1*Δ) and negative (*opt1*Δ) controls; * P < 0.05, **** P < 0.0001. A Student’s t-test using a Gaussian distribution was used to compare phagocytic indices of KN99a and *ham1*Δa, as they are both in the *MAT*a background.

### 2.7 *ham1*Δα exhibits defects in capsule attachment and transfer

First, we looked at capsule attachment. After inducing capsule formation, we subjected WT, *ham1*Δα and a known capsule mutant (*pbx1*Δ) to sonication (Fig. 7A). We found that our *ham1*Δα mutant lost significantly more capsule than WT cells, suggesting a defective attachment to the cell wall (Fig. 7B, Supplemental Fig. 2). The next avenue we investigated was capsule transfer. We incubated the acapsular mutant *cap59*Δ with conditioned media from WT, *ham1*Δα, *pbx1*Δ and *cap59*Δ to see how well the capsular material could attach to the cell wall of *cap59*Δ (Fig. 7C). As expected, our *cap59*Δ mutant could not transfer capsule at any dilution tested due to the lack of capsule. Our *pbx1*Δ mutant also was unable to transfer capsule at dilutions higher than 1:5 which was unsurprising due to its known capsule defects (22, 23). However, our *ham1*Δα also struggled to transfer the capsule at higher dilutions >1:750, whereas the WT capsule had no problem at these dilutions (Fig. 7C, Supplemental Fig. 2). This indicates at least a small defect in either structure or reduced release of capsule. Defects indicated by sonication and transfer in our *ham1*Δα prompted us to look at shedding and biofilm production, which could further confirm issues with capsular release (24).

**Figure 7:**
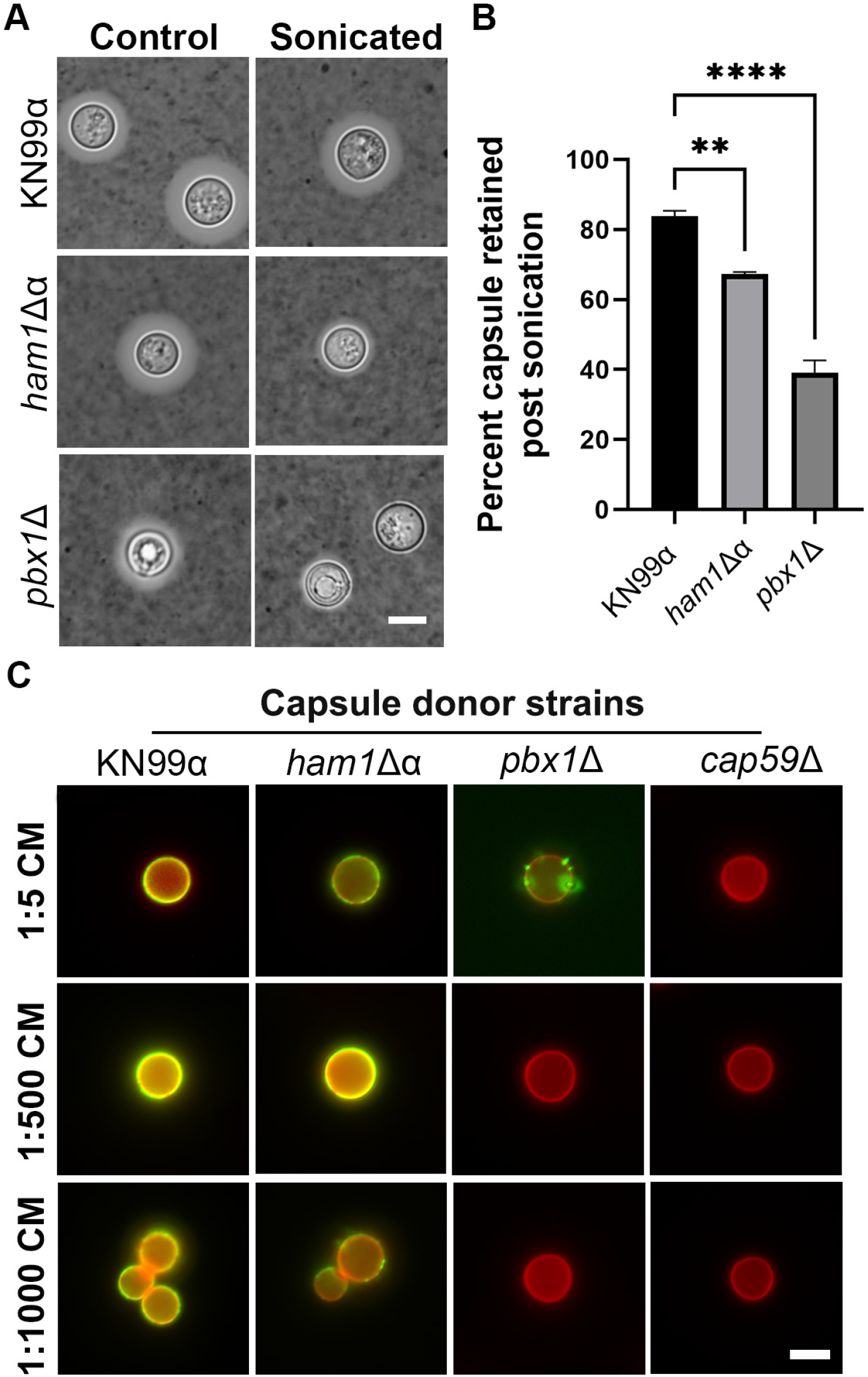
*ham1*Δα exhibits defects in capsule attachment and transfer. (A) Representative images of WT, *ham1*Δα and *pbx1*Δ capsules stained in India Ink before and after sonication. Scale bar represents 5μm. (B) Quantification of the percentage of capsule retained post sonication. One-way ANOVA with a Dunnet’s multiple comparison test relative to WT; ** P < 0.01, **** P < 0.0001, N = 20 cells for all strains. (C) Representative images of capsule transfer assay; capsule visualized with Alexa488-conjugated 3C2 antibody and cell wall with Calcoflour white (CFW). CM, conditioned media. Scale bar is 10μm.

### 2.8 *ham1Δ*α has altered capsule shedding in different medias and higher biofilm production

To investigate the possibility of defects with capsule release, we assessed the ability of *ham1*Δα to shed capsule in different media by electrophoresis and immunoblotting (24, 25)(Fig. 8A). We observed specific differences in the shed polymer size in nutrient-rich YPD medium with potential differences in size and amount in the capsule inducing medium DMEM and the minimal growth medium YNB (Fig. 8A). To get a more quantitative measurement of amounts sheds, we measured GXM concentration by ELISA (24). In YPD we saw a significant increase in the amount of capsule shed in our *ham1*Δα (Fig. 8B, Supplemental Fig. 2). We saw no significant difference in the amount of capsule material shed in YNB or DMEM relative to WT (Fig. 8. B-D, Supplemental Fig. 2). For all media, our capsule shedding mutant *pbx1*Δ shed less than WT, as expected, and our a capsular mutant *cap59*Δ did not shed any capsule. Given the association between capsule and biofilm formation, we next assessed how well *ham1*Δα could produce a biofilm using the XTT biofilm assay (24). We found that our *ham1*Δα produced larger biofilms than our WT as did *pbx1*Δ, whereas no biofilm was produced in the acapsular mutant *cap59Δ* (Fig. 8E).

**Figure 8:**
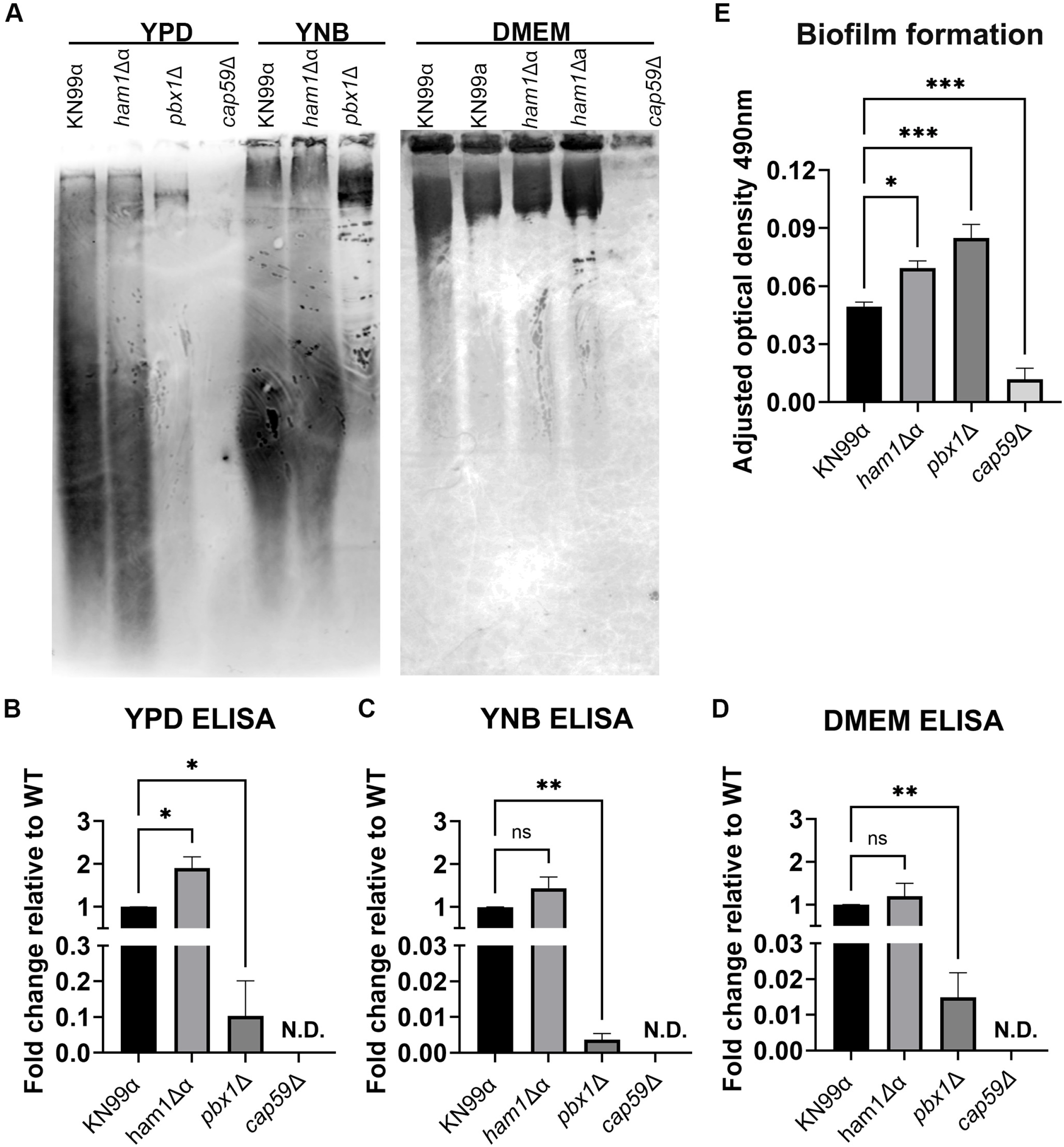
*ham1*Δα exhibits increased exopolysaccharide shedding and biofilm production. (A) Representative GXM immunoblots of exopolysaccharide shedding in capsule non-inducing media (YPD and YNB) and in the capsule inducing medium DMEM. (B - D) Quantification of shed exopolysaccharide in YPD (B), YNB (C), and DMEM (D) via Sandwich ELISA, shown as fold change relative to WT. N.D., not detected. One-way ANOVA with Dunnet’s multiple comparison test relative to WT; * P < 0.05, ** P < 0.01; N = 3 for all ELISAs. (E) Biofilm formation was quantified by the XTT reduction assay, measured by absorbance at 490nm. The *cap59*Δ is known to not form biofilms and is used as the negative control. A one-way ANOVA with multiple comparisons was run relative to WT (KN99α); * P < 0.05; *** P < 0.001; N = 3 biological replicates.

### 2.9 *ham1Δ*α shows no difference in virulence capacity in *G. mellonella*

Lastly, to investigate whether the role of *HAM1* in mating and proper capsule function translates to virulence defects, we performed infections using the invertebrate model system *Galleria mellonella* (26). We infected G. *mellonella* with 1×10^5^ log phase cells/larvae and, surprisingly, saw no significant difference in survival between WT and *ham1*Δα infected larvae (Fig. 9A, Supplemental Fig. 2). On the other hand, the *pbx1*Δ mutant showed attenuated virulence which is consistent with previous studies (22). We also assessed fungal burden at time of death, but saw no significant difference in the fungal burden of any of the strains (Fig. 9B, Supplemental Fig. 2).

**Figure 9:**
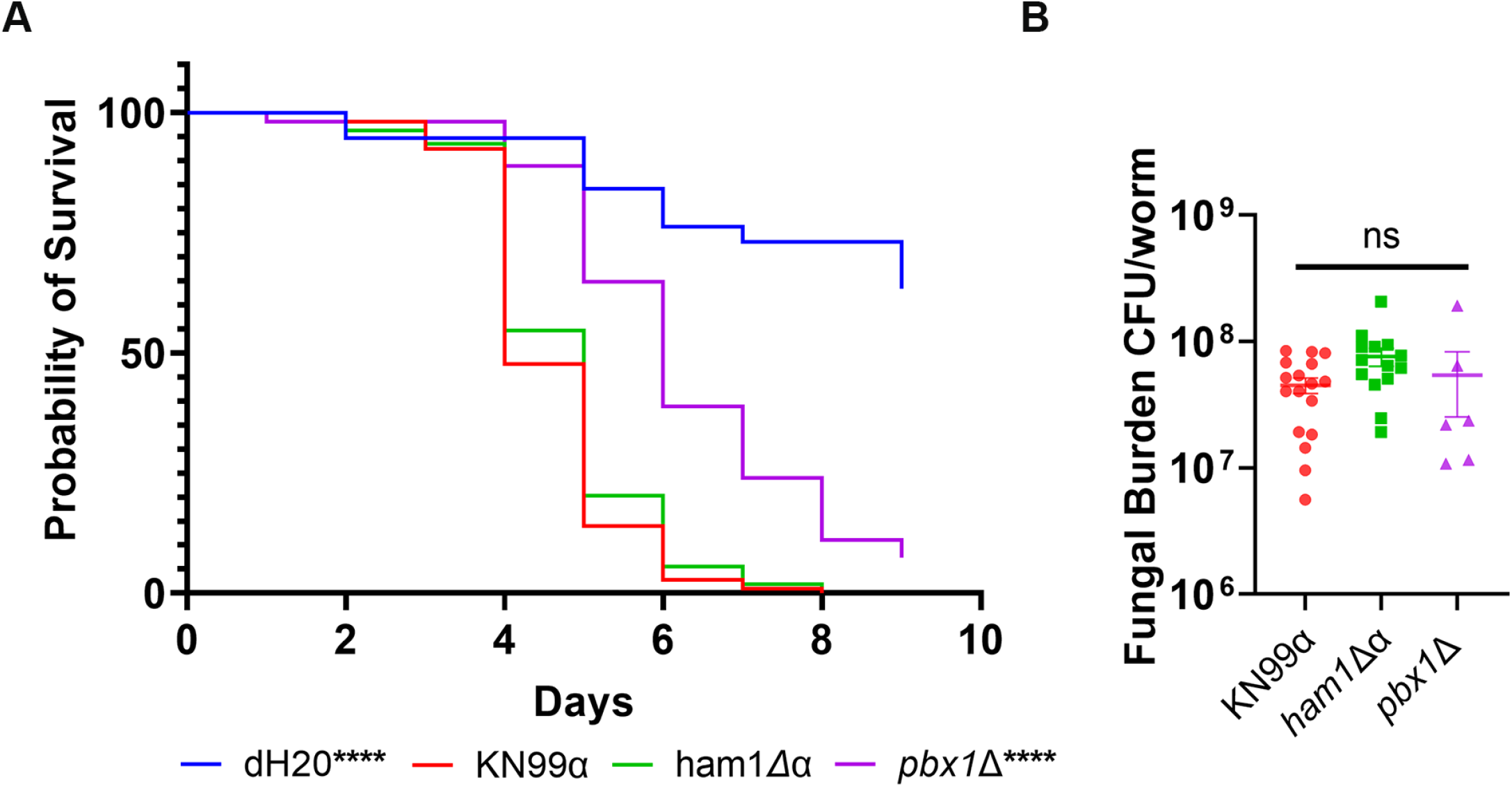
*ham1*Δα is as virulent as WT in *G. mellonella* infections. (A) Kaplan-Meier survival curve of KN99α (n=107), *ham1*Δα (n=108), *pbx1*Δ (n=54), and dH_2_O (n=68). Significance determined by Mantel-Cox test, **** P < 0.0001. (B) Fungal burden assessment of infected worms at time of death. One-way ANOVA with a Dunnet’s multiple comparisons test relative to WT; KN99α N = 17, *ham1*Δα N = 15, and *pbx1*Δ N = 6.

## 3. Discussion

Mating is a vital part of the lifecycle of the *C. neoformans* species complex. It allows for sexual reproduction, increasing the genetic diversity among progeny and overall propagation of the species. Despite having a well-defined sexual cycle both in unisexual and bisexual scenarios, not all the important regulators of mating are known, especially in *C. neoformans* (serotype A strains). Our results presented here are consistent with *HAM1* acting as a negative regulator of mating specifically in *C. neoformans*, with additional direct or indirect roles pertaining to the capsule. Moreover, the mating-associated morphological changes have been suggested to impact infection in multiple ways (13, 14), hence understanding how it is regulated may yield novel and more effective ways to prevent, or treat, this disease.

Given the proposed role of HAM-13 in *N. crassa*, we focused on the ability of cryptococcal *ham1*Δ mutants to fuse and produce recombinant progeny. Throughout our investigations, *ham1*Δ mutants consistently outperformed the WT strain in hyphal formation and production of progeny. This suggests that *HAM1*, like its homolog in *N. crassa*, is somehow connected to cell-to-cell communication and fusion of fungal cells. However, these phenotypes were only present in *C. neoformans* and not in *C. deneoformans*, precluding us from interrogating more in-depth the potential signaling pathways affected by *HAM1*. Mating studies in *Cryptococcus* have been mostly done in serotype D strains (*C. deneoformans*) rather than in serotype A, as in this study, because of the natural propensity for mating in *C. deneoformans*. Hence, although surprising, it is not unexpected that *HAM1* deletion yields different phenotypes between the sister species. Additionally, we found the *HAM1* effects in *C. neoformans* were exclusively found in the V8 mating-inducing medium as there was no significant increase in the amount of filamentation observed in either Musharige-Skoog (MS) or Filamentation Agar (data not shown). V8 media is considered the “gold standard” for looking at mating interactions in *C. neoformans*, while the others are mostly used in *C. deneoformans* (27), potentially explaining why we only see phenotypes in V8. Given that mating in *C. neoformans* is less well understood, and considering the larger epidemiological impact of serotype A strains which comprise 95% of clinical isolates and have been associated with higher virulence potential (28, 29), we chose to remain in the serotype A strain for these mating assessments. Still, as discussed below, prior transcriptomic data published for *C. deneoformans* shows *HAM1* levels are dramatically downregulated under mating conditions (30), relative to YPD, similar to our findings in *C. neoformans*.

The potential role of *HAM1* in filamentation was very apparent in our *ham1Δ* cellular fusion assay, where colonies exhibited a dry desiccated form which were in fact aggregates of hyphae. However, when these dry colonies were re-streaked, they returned to a “normal” colony morphology. The presence of the filamentous form in the absence of any mating pressure shows that the progeny of our *ham1*Δ strains have an even stronger drive towards the mating form but that over time, the signaling to return to a normal yeast state is reestablished. This, however, could impact the fitness of these strains under non-mating conditions. Thus, we were surprised to find that our *ham1Δ*α has a fitness disadvantage in both mating permissive and non-mating conditions, whereas our *ham1*Δa performed better in mating permissive and non-mating conditions. One of the main differences between cellular competition and other classical mating assays in *C. neoformans* is that in the latter, opposite mating type partners are in very close proximity which masks issues with searching for mating partners. In our *ham1*Δα competitions under mating conditions, *ham1*Δα while good at making hyphae and producing progeny with a partners in close proximity, may lack the ability to effectively forage for mating partners in nonstatic conditions. The disadvantage in non-mating conditions may be the result of two possibilities. First, *ham1*Δα has a drive to mate that outweighs the drive to cellularly divide in the presence of an opposite mating partner. Second, in *S. cerevisiae* prolonged exposure to mating pheromones results in programmed cell death (31). This same principle may apply in this case in that the presence of an opposite mating partner and the generation of *MAT*α pheromone may induce inhibition of cellular growth or death in *ham1*Δα cells. Surprisingly, in our *ham1*Δa competition, we observed an increase in competitive fitness in both non-mating and mating conditions. These observations may be the result of the differences in the morphological changes happening to the different mating types. For example, in *C. deneoformans* the *MAT*α cell responds to pheromone by forming conjugation tubes while the *MAT*a cells become enlarged (32). To the best of our knowledge, these pre-cellular fusion structures have been observed only in *C. deneoformans* and are defined by their *MAT* loci (33-35). If this pre-cellular fusion morphology occurs in *C. neoformans*, it’s possible that in the mating permissive competitive scenario with the *ham1*Δa strain, the WT *MAT*α mating partner is able to produce conjugation tubes and effectively seek the *MAT*a mutant partner. This would not happen with the *ham1*Δα strain seeking the WT *MAT*a partner, explaining the different results. This also highlights the possibility that *HAM1* may have a greater effect on *MAT*α strains relative to *MAT*a ones. This would not be a new concept. A previous study looking at the global transcriptional repressor *TUP1* found mating-type specific phenotypes (36). Disruption of *TUP1* led to different gene expression profiles in *MAT*α vs *MAT*a despite having very similar cellular fusion and filamentation phenotypes. This finding implies that even amongst lab strains there is heterogeneity with respect to competitive fitness depending on the *MAT* locus, and that not all WT strains will behave the same when in a competitive environment.

We wondered if *HAM1* was acting on the pheromone MAP kinase pathway, but we were unable to obtain clear results from the qRT-PCR. This was not surprising given that serotype A strains do not respond to mating signaling uniformly or at the same time, hence the RNA samples represent a mixture of states where mating cells are a minority. This was the case when measuring *MAT*α, *MAT*a, *PUM1*, and *CFL1* during unilateral and bilateral crosses, where we saw no statistically significant differences relative to WT crosses. Still, we saw a consistent and significant repression of *HAM1* on V8 early on, as expected if acting as a negative regulator of mating. Notably, repression of *HAM1* would occur in V8 only in the presence of both mating partners, since in monocultures of *MAT*α or *MAT*a strains, *HAM1* was induced in V8 relative to YPD, again consistent with it being a negative regulator of mating. Interestingly, in *mat2*Δ monocultures in V8, we still saw an induction of *HAM1*, whereas in *cpk1*Δ monocultures, missing Mat2’s upstream kinase, *HAM1* was repressed. This is consistent with a model where upon pheromone pathway activation, *HAM1* is repressed in a Cpk1-dependent but Mat2-independent manner. However, in the absence of pheromone activation, *HAM1* is induced, indicating additional regulatory inputs into *HAM1*. While mating is predominantly controlled by the pheromone response pathway, previous studies have shown that calcineurin signaling is essential for mating, haploid fruiting and spore production (37, 38). Hence, we tested a potential involvement of *HAM1* in that pathway by assessing sensitivity to calcium stress, but found no defects with our *ham1*Δ mutants (Supplemental Fig. 3). Still, this does not rule out the possibility that calcineurin impinges on *HAM1* regulation. In fact, a study looking at the calcineurin-regulated genes necessary for thermotolerance found *HAM1*. Upon shift from 24°C to 37°C, there is a 14-17-fold increase in *HAM1* levels, while in the *cna1*Δ is only 8-fold (38).

One of the hallmarks of mating in *C. neoformans* is the morphological transition from yeast to hyphae. The hyphal form of *C. neoformans* has been shown to be unable to cause disease in a mammalian host in part because it induces an immune response shift from a disease-permissive TH-2 to a disease-protective TH-1 proinflammatory response. When examining previously published RNA-seq data, we noted that *HAM1* transcripts were some of the highest induced in cryptococcosis patients (39, 40). This finding is intriguing for two reasons. First, it lends support to the idea that to prevent a morphological transition in this low nutrient environment negative regulators of mating would be highly expressed. Secondly, it may indicate additional roles for *HAM1* directly related to maintenance of virulence factors providing further evidence of the link between mating and virulence. However, our initial survey of the key virulence mechanisms did not show any obvious defects in *ham1Δ* mutants. Still, we decided to take a closer look at the capsule since, as a downstream target of Pfa4, it’s possible that *ham1*Δ may share defects related to the cell wall or capsule as exhibited by the *pfa4*Δ mutant. Additionally, since one of the major phenotypes in the *ham1Δ* relates to filamentation, we wanted to investigate if there was a possible cross over between the cellular circuits governing morphology and capsule production and maintenance.

Although capsule induction was similar, *ham1*Δ mutants exhibited a defect in capsule attachment and shedding. This was similar to phenotypes exhibited by *pbx1*Δ and *pbx2*Δ strains, both mutants with abnormal capsule structure (22, 23). While defects were milder than *pbx1Δ,* capsule attachment and transfer were reduced in the *ham1Δ.* Capsular shedding, however, was increased in YPD, a capsule non-inducing medium, and trended towards higher shedding in capsule-inducing media. This suggested a level of involvement of *HAM1* in proper capsule formation that could be a direct or an indirect consequence of dysregulated mating signaling. Capsule regulation and structure during mating conditions is a topic that has received little attention. It is reasonable to envision changes in capsule accompanying the morphological changes present during mating. Hence, these capsule defects seen in *ham1*Δ mutants might be a consequence of the abnormal mating signaling occurring in the absence of *HAM1*. Consistently, normal capsule production is required for biofilm formation. Biofilm production, while not seen as a virulence factor for *C. neoformans*, was likely developed as a survival tactic against environmental predation (41). The *ham1*Δα mutants have altered biofilm formation, again suggesting defects in capsule attachment and altered capsule structure. With all the similarities in capsule defects shared between *pbx1*Δ and *ham1*Δα, we next wanted to assess the virulence capabilities of *ham1*Δ.

Deletion of *pbx1Δ* results in attenuated virulence in a mouse model so we expected to see a similar defect in our *ham1*Δα (22). To our surprise, there were no significant defects in virulence potential using the *G. mellonella* infection model relative to WT (median death rate: WT, 4 days; *ham1*Δα, 5 days; and *pbx1*Δ, 6 days). Likewise, there were no differences in fungal burden, even in the *pbx1*Δ-infected larvae. This could be attributed to the fact that they lived longer, as the burden was assessed at time of death. More detailed virulence assays will be needed to confirm if there are defects in *ham1*Δ mutants affecting its pathogenesis.

Taking all of these findings together we propose a model where *HAM1* negatively regulates the mating cycle of *C. neoformans* (Fig. 10). We envision two possible locations for where it could be acting: first, it could act directly on pheromone activated genes, inhibiting their function; or secondly, it could be directly involved in the yeast to hyphal transition. Although we do not have evidence for it, it is possible it is acting at later stages as an inhibitor of recombination or spore generation, although it is unlikely given that we see *HAM1* expression changes only at day 1 under mating conditions. Regardless, by acting as a negative regulator of the mating cycle we believe it prevents mating interactions from occurring during pathogenic conditions. This could be a potential reason for why *HAM1* is so highly expressed in the CSF of cryptococcosis patients. CSF is a very nutrient limited environment which may provide the right pressures for cells to undergo a mating interaction, but that would be detrimental to pathogenesis. The signal to activate *HAM1* under non-mating conditions, however, remains unclear.

**Figure 10:**
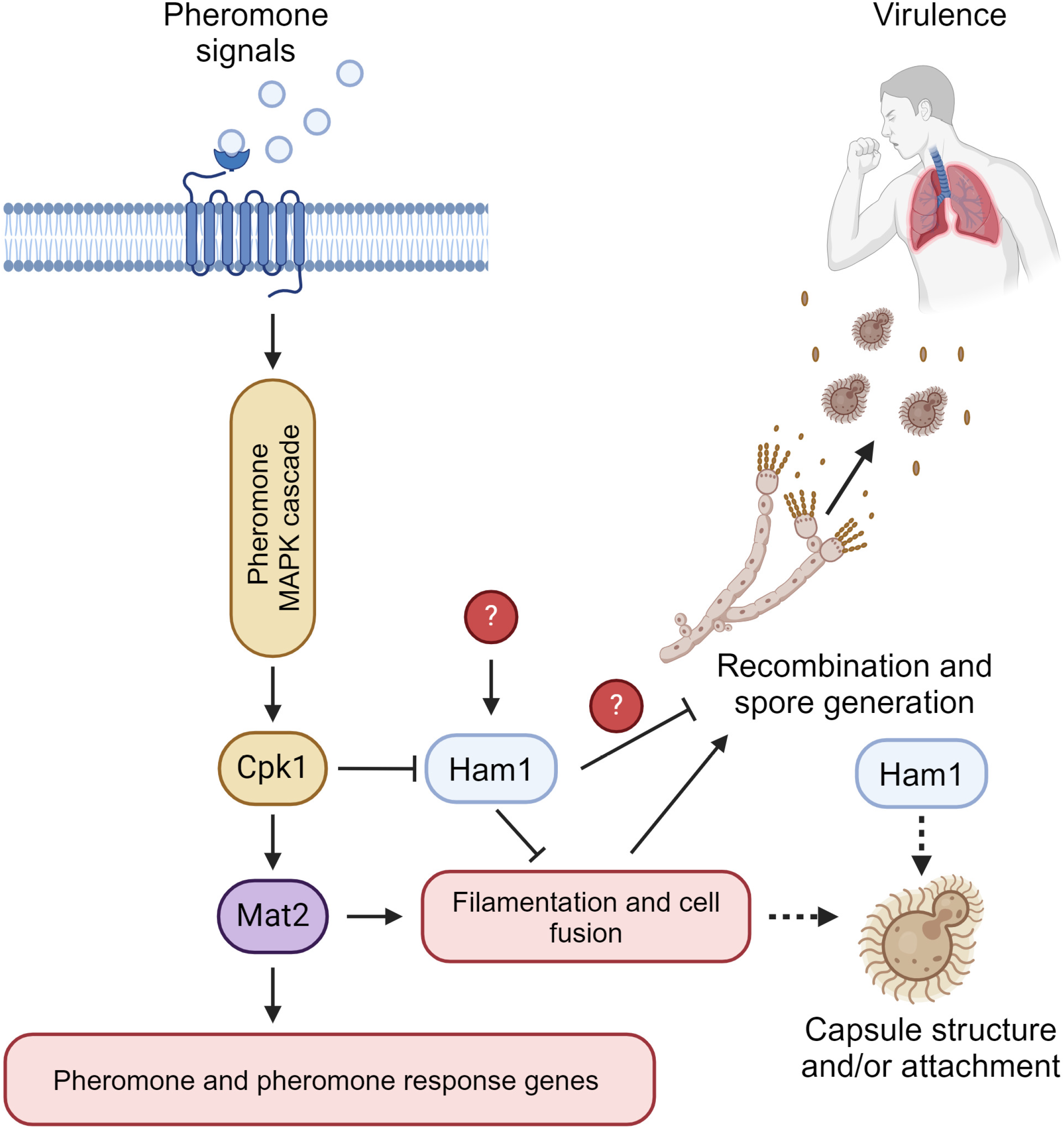
Proposed model of HAM1 involvement in C. neoformans biology. Based on this study, we believe that *HAM1* acts as a negative regulator in the mating cycle of *C. neoformans*. Upon activation of the pheromone response pathway, *HAM1* is repressed in a Cpk1-dependent manner. However, under non-mating conditions, *HAM1* is induced by unknown signals. When induced, *HAM1* regulates mating by inhibiting pheromone and pheromone response genes directly, or somehow inhibiting hyphal formation directly. During pathogenic conditions, we believe that *HAM1* is highly expressed to prevent mating from occurring due to the low nutrient environment of a mammalian host. The changes we see in capsule could be a direct effect of *HAM1*, or an indirect result of altered mating signaling in the absence of *HAM1* (dashed lines). This figure was created with BioRender.com.

By uncovering a role for *HAM1* in *C. neoformans* biology we have shown another potential intersection point between virulence and mating. Both processes are vital to the propagation and survival of the species and the presence of regulators with roles in multiple pathways reflects just how important these are. Investigation of mating regulators not only improves our knowledge of how the system works but also opens new avenues of investigation for therapeutic interventions. Through a better understanding of fungal mating, we may be able to find more intersection points between virulence and mating pathways to fully understand how these two processes are connected. We hope this study serves to raise awareness about the importance of mating for virulence and stirs more investigations into this phenomenon.

## 4. Methods

### Strains and growth materials

All serotype A strains were in the KN99 background (Supplementary Table 1). All *ham1*Δ strains and *HAM1*-FLAG tagged strain were generated using biolistic delivery (see methods below and Supplementary Table 1). V8 agar medium was prepared using original recipe V8 juice, bacteriological grade agar and dH20. V8 medium was adjusted to the appropriate pH (5 for serotype A, 7 for serotype D) using 1M hydrochloric acid or sodium hydroxide tablets, respectively. 25mM CuSO_4_ was added after autoclaving before pouring plates.

### Phenotyping

Strains were grown overnight in 5mL of YPD liquid culture diluted to an OD_600_ of 0.2 and left to grow for 2 doublings. Strains were washed in 1x PBS and counted using a Bio-Rad TC20 automatic cell counter and diluted to final concentration of 1×10^7^ cells/mL. Cells were then serially diluted by 10-fold to a final concentration of 1×10^3^ cells/mL. 5μL of culture was spotted onto the following plates: YPD, YNB, YPD with 1M NaCl, 0.5 mg/mL Caffeine, 0.1- 0.3% SDS, 0.35-0.5M CaCl_2_, 0.5 mg/mL Calcofluor White (CFW) Oxidative (H_2_O_2_) and Nitrosative Stress (NaNO_2_). Plates supplemented with stressors were incubated at 30C for 2-3 days. Plates that had no stressors were incubated at 30°C and 37°C to assess temperature defects. All plates were imaged on the Bio-Rad EZ Gel Doc Imager.

### Fungal Genomic Manipulation

We generated all *ham1*Δ mutants using the split marker method to delete the entire *HAM1* coding region. Fragments were delivered to candidate KN99α and KN99a cultures via biolistic delivery (Bio-Rad PDS-1000). Deletion candidates (from at least two independent transformations) were assessed using PCR as well as mating phenotype analysis to confirm deletions. Two of these independent deletion mutants were tested in all mating assays. To perform immunoprecipitations needed for the click chemistry analyses, a FLAG-tag strain was generated using the 4x FLAG-Tag plasmid to tag the C-terminal end of Ham1. This fragment with overlapping homology to the *HAM1* coding sequence was synthesized by Genewiz and again delivered to candidate cells via biolistic delivery. This *HAM1*-FLAG tagged strain was functional as it was indistinguishable from WT in mating and capsule assays.

In addition to generating deletion mutants via biolistics, we generated two independent *ham1*Δa candidates via mating of KN99a with *ham1*Δα (Supplementary Table 1). Strains were grown in 5 mL of YPD for 48 hours, spun down and washed twice in dH20, counted, and normalized to a concentration of 1.5 x10^7^ cells/mL. Equal cell volumes of each mating pairs were mixed and incubated at RT for an hour. 100-200 μL of this mixture was spotted onto V8 mating plates and incubated in the dark at RT for 7 days. Filaments from each mating interaction were scraped using a sterile toothpick and resuspended in dH20. Spores were plated on YPD with selection agar plates and streaked twice with selection for single colonies. Isolated single colonies were tested for mating type using a mating interaction with known tester strains (JEC20 and JEC21). These *ham1*Δa mutants behaved similar to the mutants generated by biolistics. We also generated a *ham1*Δα with the NEO cassette by CRISPR (Supplementary Table 1) using the method published by the Madhani lab (42).

Lastly, because we were unable to generate a complemented strain on the same loci by biolistics, we tested a commercially available *ham1*Δ deletion which we refer as C07 (the well number on the deletion collection plate). All 3 independent mutants showed the same phenotypes (Supplementary Fig. 2 and data not shown). We also used a plasmid complementation approach by cloning the *HAM1* locus (including its native promoter and terminator) into plasmid pMSC042-Neo (Supplementary Fig. 4A) and electroporating into *ham1*Δα-NAT. We used this strain to asses complementation of the capsule attachment phenotype (Supplementary Fig. 4B) and the biofilm formation alteration (Supplementary Fig. 4C).

### Cell preparation, protein extraction, and immunoprecipitation for click-chemistry

Using “clickable” fatty acid probes, such as alkynyl palmitate (alk15; Click Chemistry Tools #1165), that are metabolically incorporated into normal substrates by the cell, we can then carry out a bioorthogonal Cu(I)-catalyzed Huisgen 1,3-dipolar cycloaddition reaction (the click-reaction) to conjugate the fatty acid-modified proteins to easily measurable reporters such as fluorophores or affinity tags. To confirm Ham1 is palmitoylated, *HAM1*-FLAG and KN99a strains were grown in 50 mL of YNB medium, pH 7.0, and once an OD of 0.5 was reached, strains were fed with 15 μL of 5 μM alk15 and incubated for 1hr at 30°C with shaking. Cells were washed twice in cold dH_2_0 and flash frozen until ready for use. Cells were thawed on ice and disrupted in lysis buffer (100 mM Tris-HCl pH 7.4, 0.5% Triton X-100, 150 mM NaCl, 13% w/v glycerol, protease inhibitor tablet, Roche) by bead beating (mini bead beater-16, Biospec product #607) for 5 cycles of 1 minute on, 1 minute rest on ice. Cell extracts were spun down at 13,000 x g for 10 minutes at 4C, and protein concentration in the supernatant determined by a detergent compatible Bradford assay (Thermo Scientific #23246).

Based on described methods (43), 1μg of protein lysates from fed and unfed cell populations were incubated with 100 μL of 50% slurry of anti-FLAG M2 affinity gel (A2220; Sigma-Aldrich) for 20 minutes on a nutator at 4°C. The protein-bead mixtures were then loaded onto a gravity column (TALON 2 mL Disposable Gravity Column, Takara #635606). Lysate was loaded into the column 5 times with gravity flow and then washed in a 1M NaCl buffer 3 times. Beads were then washed twice with 150 mM NaCl buffer before resuspended in Click Chemistry buffer according to the manufacturer’s protocol (#1262, Click Chemistry Tools) apart from the AFDye 680 Azide Plus reporter (Click Chemistry Tools #1512) which was added at a concentration of 1.25uM. Excess reactants were removed by spinning down anti-FLAG M2 affinity gel and washing twice in 1x PBS. Beads were then resuspended in 1x Laemmli Buffer and heated at 70°C for 10 minutes. After SDS-PAGE the 10% polyacrylamide gel was destained in Click Chemistry Destain (40% methanol, 50% acetic acid, 10% water) overnight and gel visualized on an Odyssey Imaging System (LI-COR Biosciences).

### Cellular Fusion

Cellular fusion efficiency was assessed using overnight cultures grown in YPD that were washed twice in dH20. Cultures were normalized to an OD of 0.5 and resuspended in dH20. Each cross is comprised of cells of opposite mating type and different antibiotic resistance cassettes to determine the fusion efficiency of each cross. Equal volumes of each strain are mixed for their respective crosses and spotted in 50 μL increments on V8 mating agar. Each cross is spotted in triplicate and left to dry for 20 minutes. Once all plates were dry, they were transferred to a mating chamber (DM1, Mycolabs) and left in the dark for 5 days at ambient room temperature with a relative humidity of 40-50%. After incubation, each cross was scraped and resuspended in 1 mL of dH20 and normalized via cell count to 1×10^7^ cells/mL. For quantification, 100 μL of undiluted, or 10 and 100-fold dilutions of the 1×10^7^ cells/mL suspension, was plated on both double and single antibiotic YPD resistance plates (NAT and NEO). Once plates were dry, they were transferred to a 30°C room, incubated for three to four days, and counted for CFUs.

### Hyphal staining

Single colonies from mutant crosses were scraped and resuspended in dH20. Colonies were then stained with either 5 μg/mL DAPI or 100 μg/mL Calcofluor White (CFW) and incubated at 30°C in the dark for 20 minutes. Cells were then washed three times in 1x PBS and resuspended in 100 μL. Cells were then imaged using a 100x oil immersion objective in a Zeiss Axio Observer microscope.

### Assessment of Bisexual Mating

Adapted from bisexual mating protocol from Sun et al., 2019 (27). Briefly, strains were grown overnight in YPD, washed twice in dH20 and normalized to an OD of 1. Cells of opposite mating type for each cross were mixed in equal amounts in a separate tube and subsequently spotted in 20 μL amounts onto V8 agar plates. All crosses were spotted in triplicate and left to dry for 20 minutes. Once all plates were dry, they were transferred to a mating chamber (DM1, Mycolabs) and left in the dark for up to 4 weeks with a relative humidity of 40-50%.

### Cellular Competition Assessment

Each cross consists of three strains: two of the same mating type but with opposing antibiotic resistance cassettes (competitor strains), and one of the opposite mating type with no antibiotic resistance (donor strain). Overnight cultures for each strain were grown in YPD until log phase. Cultures were then washed twice in dH20, counted, and normalized to a concentration of 1×10^5^ cells/ml. Competitor and recipient strains were mixed in equal amounts for each cross. 30 μL of each cross was plated in triplicate on V8 and YPD medium and left to dry for 20 minutes. Once all plates were dry, they were transferred to a mating chamber and left in the dark for 10 days with a relative humidity of 40-50%. After the 10 days crosses were scraped and resuspended in dH20. Crosses in YPD were diluted 100,000x and crosses in V8 were diluted 10,000x. 100 μL of the respective dilutions was plated in single antibiotic supplemented YPD plates (NAT or G418). Plates were incubated for 2-3 days and imaged for CFUs counting using a Bio-Rad Gel Doc EZ imager.

### RNA isolation from bisexual mating crosses

Bisexual mating crosses are set up as previously described and allowed to incubate for 1, 3, 5 and 7 days. Cells are then harvested, washed once in dH20, washed with RNA stop solution and flash frozen and stored at -80°C until ready for extraction. A TRIzol-extraction (Life Technologies #15596-026) protocol is used followed by an RNA clean up step using the ZYMO RNA Clean and Concentrator kit (cat # R1019). Up to 1ug of RNA is then converted into cDNA using the NEB LunaScript® RT Supermix Kit (NEB #E3010) followed by qPCR using the NEB Luna Universal qPCR Master Mix (NEB #M3003). The ΔΔ_CT_ method was used for quantification of relative transcript level normalized to YPD samples (44).

### Capsule induction and visualization with India Ink

Cultures were grown overnight in 5 mL of YPD medium. Cultures were spun down at 3,000 x g for 5 minutes and resuspended in 1 mL of PBS. After cell count, 1×10^7^ cells/ml were transferred to a new microtube and pelleted. Pellet was resuspended in 1 mL of Dulbelcco’s Modified Eagle’s Medium (DMEM) (Corning; VWR 45000-304). This suspension was diluted 10-fold by adding 500 μL to 4.5 mL of DMEM to a final concentration of 1×10^6^ cells/mL. 1 mL aliquots of this cell suspension were added into a 24 well culture plate and incubated at 37°C with 5% CO_2_ for 24 hours. After 24-hour incubation, samples were transferred to microtubes and spun down at 3,000 x g for 5 minutes. Samples were washed in PBS and resuspended in a final volume of 50 μL. In a separate tube, 10 μL of sample were mix wth India Ink and a drop of the mixed sample and India ink was pipetted onto a Polylysine-coated slide and imaged on a Zeiss Axio Observer Microscope.

### Capsule Sonication

Capsule is induced as described above and cells were resuspended in 1 mL of 1x PBS. A portion of the cells was aliquoted (unsonicated controls) and the rest were subjected to sonication using a tip sonicator (Branson #SFX250) at 20% power with 0.5 pulse for 20s and kept on ice. Once all strains had been sonicated, all samples were mixed with India ink as described above and visualized on a Zeiss Axio Observer Microscope at 100x oil immersion magnification. The capsule radius was measured using FIJI (45).

### Capsule Transfer

Capsule transfer assays were performed as previously described (23). Capsule donor strains were grown in 5 mL of YPD liquid culture for 5 days prior to the experiment. On day 4 a *cap59Δ* acceptor culture was grown overnight. Donor cultures were normalized to cell counts and spun down for 5 minutes at 3,000 x g and the top 1.5 mL was filtered through a 0.22μm filter (Avantor #76479-024). 1 mL from this filtration step was heated at 70°C to create conditioned media. Acceptor cells were washed twice with 1x PBS and counted on a Bio-Rad TC20 automatic cell counter and diluted to final concentration of 5×10^6^ cells/mL in a volume of 400μL. Conditioned media was added in the appropriate dilution to acceptor cells and incubated for 1hr with gentle rotation. Cells were pelleted at 3000 x g for 5 minutes and washed twice with 1x PBS. Resuspend cells in a volume of 100 μL and added Cy3 conjugated 3C2 anti GXM antibody to a final concentration of 8ug/ml. Incubated for 1 hour with gentle rotation. Cells were pelleted at 3000 x g for 5 minutes and washed twice with 1x PBS. Cells were resuspended in 100 μL and 100 ug/mL concentration of CFW was added and incubated for 20-30 mins with gentle rotation. Cells were pelleted at 3000 x g for 5 minutes and washed twice with 1x PBS. Cells were resuspended in a final volume of 50 μL of 1x PBS.

### Capsule Shedding via GXM Immunoblotting

GXM immunoblotting assays were performed as previously described (24, 25). Briefly, cells were inoculated into YPD and allowed to grow for 24 hours. Subsequently dilutions of 1:100 were performed and cells were inoculated into the desired media of interest with the exception of DMEM where 1×10^6^ cells/mL were added to 5 mL of fresh DMEM. Cells were then incubated in their respective medias for 3 days at 30°C with shaking at 225 rpm. DMEM cultures were incubated at 37°C with 5% CO_2_ to induce capsule formation and subsequent shedding. After 3 days, the cultures were quantified on a cell counter (Bio-Rad TC20), and the whole culture were spun down for 5 minutes at 3000 x g. The top 1.5 mL was filtered through a 0.22μm filter. 1 mL from this filtration step was heated at 70°C for 10 minutes to create conditioned media. Subsequent conditioned media was loaded onto a 0.6% Megabase agarose gel (Bio-Rad #1613108) and run for 8-10 hr at 40V. To normalize the amount of GXM loaded onto the gel, cell counts were taken for each strain in each media. The amount of GXM loaded was normalized to the lowest cell count in a total volume of 100 ul. When the dye front was at the bottom of the gel, the electrophoresis was stopped and the gel used to assemble the immunoblot sandwich. Using a Nylon membrane, the sandwich was assembled as follows, starting from the bottom: wick, 3 pieces of thick blotting paper, gel, membrane, 3 pieces of blotting paper, and paper towels (approx. 5 cm in height). The immunoblot sandwich was incubated with a 20x Sodium Citrate buffer for 10-12 hours or until all the paper towels were saturated. The membrane was blocked for 48 hours in 1× TBS-5% milk and incubated for 1 h in 1×TBST-1% milk with 1 µg/mL of anti-GXM monoclonal antibodies F12D2 and 1255. The membrane was rinsed three times in 1× TBST and incubated for 1 h in 1× TBST-1% milk with Odyssey antibody at 1:10,000. The membrane was rinsed again three times in 1× TBST and imaged on the Odyssey Imaging System (LI-COR Biosciences).

### Capsule Shedding via ELISA

GXM ELISA was done as described using a kit by IMMY-Labs (#CRY 101)(24). Conditioned media from cells was obtained as described above for GXM immunoblots. Conditioned media was diluted by a factor of 1×10^-4^ or 1×10^-6^ using serial dilutions due to the sensitivity of the assay. Lyophilized GXM was provided by the Brown Lab at the University of Utah and diluted to a concentration of 2mg/mL and working stocks were generated. The highest standard used in the ELISA was 78 ng/mL.

### XTT Biofilm Assay

Assay was performed as previously described (46). Strains were grown overnight in YPD, cells were then counted (Bio-Rad TC20) and diluted to 1×10^7^ cells/mL in DMEM, and 100 μL of all of cell suspension was seeded into wells of a microtiter plate. Plates were incubated at 37°C + 5% CO_2_ for 48 hr, washed with 1x PBS using a microtiter plate washer (405LS, Agilent), and 100 μL of XTT/menadione solution (Sigma) was added to each well. The plates were incubated for 3 hr and 75 μL of the supernatant was removed and transferred to a new microtiter plate. Plates were read in a microtiter plate reader at 490 nm.

### Uptake Assay

Uptake assays were performed as previously described (8). PMA-differentiated THP-1 cells were incubated with Lucifer Yellow-stained fungal cells that had been opsonized with 40% Human serum collected from healthy donors (approved by the Notre Dame Institutional Review Board (IRB) as a non-human subject research procedure). After incubation for 1 hour at 37°C with 5% CO_2_, plates were washed with 1x PBS using a microplate washer (405LS, Agilent). Cells were then fixed with 4% formaldehyde and stained with DAPI (Sigma) and Cell Mask (Invitrogen). NaN_3_ in PBS was added to the plates and imaged on a Zeiss Axio Observer Microscope with an automatic stage. Each well was imaged using a 3×3 grid set up and resulting images were analyzed using a Cell Profiler pipeline to determine the Phagocytic Index (PI, # of internal cells per 100 THP-1) values.

### Infections with Galleria mellonella

*G. mellonella* third instar larvae were sorted and weighed as described (47). Larvae weighing 200 mg and above were used in the following infection assay. Overnight cultures were grown and measured for an OD of 0.5-1. Cells were then washed twice in 1x PBS and counted using a cell counter. A total inoculum of 1×10^5^ cells/larvae was used in a total of 5 μL. The back larval prolegs were swabbed with 70% Ethanol and 5 μL of culture was injected into the back right proleg using a Hamilton Syringe (Hamilton, #80300). Syringes were sanitized and cleaned before and after each strain with both 70% Ethanol and dH_2_O. Survival was assessed daily. At time of death, infected *G. mellonella* larvae were placed in 3 mL of dH_2_O in a 15 mL falcon tube. Larvae were then homogenized using a tissue homogenizer (VWR #10032-336) to break down tissue. Once sufficiently homogenized, worm slurry was then diluted 10,000 or 20,000x times and plated on YPD plates supplemented with either Nourseothricin (NAT) and Ampicillin (AMP) or Gentecin (G418) and Ampicillin (AMP). Plates were left to incubate for 2-3 days and then imaged on the Bio-Rad EZ Gel Doc imager for CFU quantification.

## Acknowledgements

We would like to thank Dr. Rebecca Wingert for the use of her microscope for mating images, Dr. Patricia Champion for use of her tip sonicator, Dr. Jessica Brown and Krystal Chung for their gift of purified GXM, and Dr. Scott Briggs and Smriti Hoda for training on the *G. mellonella* infection model. We would also like to thank the members of the Santiago-Tirado lab for their help with editing and reading this manuscript. Finally, we want to acknowledge all the time and effort by Nathan Glueck in the Lin lab to delete *HAM1* in serotype D strains. This work was supported by institutional funds from the University of Notre Dame. Research in the Santiago-Tirado lab is supported by grants from the NIH (R21AI171742 and R01AI177875). We declare no conflict of interest.

## Figure Legends

Supplemental Figure 1: **Cellular fusion comparisons with or without exogenous pheromone addition.** (A) Cell fusion results from unisexual and bisexual bilateral *ham1*Δ crosses, without addition of exogenous pheromone. (B – D) Cell fusion results without and with 5µM, 7.5µM, and 10µM exogenous pheromone, in WT mating controls (B), *ham1*Δα unilateral cross (C), and *ham1*Δa unilateral cross (D). In (A) a one-way ANOVA with uncorrected multiple comparisons (unpaired t with Welch’s correction) was run; * P < 0.05; *** P < 0.001. N = 3 biological replicates per condition. All others (B – D), a one-way ANOVA with multiple comparisons (Kruskal-Wallis test) was run on all conditions, none were statistically significant. N = 3 – 5 biological replicates per condition.

Supplemental Figure 2: **C07 behaves similarly to *ham1*Δ mutants in key mating and virulence assays.** The most recent addition to the *C. neoformans* deletion collection contained a deletion mutant of *HAM1* which was well number C07 (https://www.fgsc.net/). We used the C07 mutant in a variety of assays together with our *ham1*Δ mutants made by biolistics, but for simplicity and clarity, we did not include C07 on the main figures. Here, we show the phenotypes present in C07 to provide external validation of the phenotypes we see in our *ham1*Δ mutants. (A) C07 produces hyphae at a similar speed and robustness to our *ham1*Δ mutants. The first 3 rows are the same images shown in Fig. 3. (B) C07 produces similar amounts of double resistant progeny to our *ham1*Δ mutants. A one-way ANOVA was run using multiple comparisons to the WT cross. ** p=0.0014, ***p=0.0003. N = 3 biological replicates for all conditions (C) C07 has similar defects in capsule attachment to *ham1*Δ mutants post sonication. A one-way ANOVA was run using a Dunnett’s multiple comparison test of all mutant crosses to the WT cross; ** p=0.0062 (*ham1*Δα) **, p=0.0040 (C07), ***p=0.0004. N = 20 cells for all conditions. (D - F) Quantification of capsule shedding by C07 strain in non-capsule inducing media YPD (D) and YNB (E) and in the capsule inducing medium DMEM (F). All results were compared by one-way ANOVA with a Fisher’s LSD test, * P < 0.05. N = 3 biological replicates for all ELISAs (G) C07 fails to fully transfer capsule at high dilutions greater than 1:750. Representative images of capsule transfer capsule visualized by conjugated 3C2 antibody with Alexa488 fluorescent probe and cell wall stained with calcofluor white (CFW) scale bar is 10μm. (H) C07 shows no difference in survival compared to WT. Kaplan-Meier survival curve of dH_2_O=10 KN99α N = 30, *ham1*Δα N = 30 and C07 *N =* 30. No statistical differences were seen in the mutants but the dH_2_O survived significantly longer, **** P < 0.0001 by Mantel-Cox test. (I) C07 shows no difference in fungal burden. KN99α N = 7, *ham1*Δα *N =* 7, C07 N = 11. A one-way ANOVA with a Dunnet’s multiple comparison test of all conditions to WT was run and showed no significant difference between mutants and WT.

Supplemental Figure 3: ***ham1*Δ shows no sensitivities to various cell wall and membrane stressors as well as antifungal treatments.** To rule out any issues with calcineurin signaling, cell membrane, or cell wall, we tested various conditions in YPD solid agar medium. (A) Growth of indicated strains in the presence of calcium (CaCl_2_) and sodium dodecyl sulfate (SDS) at 30°C or 37°C. (B) Growth of indicated strains to test thermal stress (YPD at 30°C and 37°C), osmotic stress (NaCl), nutrient stress (YNB), and cell wall stress signaling (Caffeine). (C) Growth of indicated strains under cell wall stress (calcofluor white, CFW), oxidative stress (H_2_O_2_) and oxidative stress (NaNO_3_). (D) Growth of indicated strains in the presence of E-test strips of common antifungals (Amphotericin B and Fluconazole).

Supplemental Figure 4: **Ectopic complementation of *ham1*Δα restores the capsule shedding and biofilm formation phenotypes.** (A) Plasmid map used for ectopic complementation of *ham1Δ*α. This plasmid (pMSC042-Neo, a gift from the Doering lab) contains telomeric sequences flanking a small piece of the kanR gene. When linearized with I-SceI, the plasmid DNA is capped by these telomeric sequences, preventing degradation. (B) Results from capsule attachment assay using sonication. Shown is 2 biological replicates combined. (C) Results from biofilm formation assay using XTT reduction as readout (measured by absorbance at 490nm). Shown is the average and SD of six technical replicates. One-way ANOVA with multiple comparisons; * P < 0.05; ** P < 0.01.

Supplemental Table 1: **Strains used in this study, information and references.**

## References

1. Hawksworth DL, Lucking R. 2017. Fungal Diversity Revisited: 2.2 to 3.8 Million Species. Microbiol Spectr 5.

2. Fisher MC, Gurr SJ, Cuomo CA, Blehert DS, Jin H, Stukenbrock EH, Stajich JE, Kahmann R, Boone C, Denning DW, Gow NAR, Klein BS, Kronstad JW, Sheppard DC, Taylor JW, Wright GD, Heitman J, Casadevall A, Cowen LE. 2020. Threats Posed by the Fungal Kingdom to Humans, Wildlife, and Agriculture. mBio 11.

3. Min K, Neiman AM, Konopka JB. 2020. Fungal Pathogens: Shape-Shifting Invaders. Trends Microbiol 28:922–933.

4. Srikanta D, Santiago-Tirado FH, Doering TL. 2014. *Cryptococcus neoformans:* historical curiosity to modern pathogen. Yeast 31:47–60.

5. Maziarz EK, Perfect JR. 2016. Cryptococcosis. Infect Dis Clin North Am 30:179–206.

6. Rajasingham R, Govender NP, Jordan A, Loyse A, Shroufi A, Denning DW, Meya DB, Chiller TM, Boulware DR. 2022. The global burden of HIV-associated cryptococcal infection in adults in 2020: a modelling analysis. Lancet Infect Dis doi:10.1016/S1473-3099(22)00499-6.

7. Fisher MC, Denning DW. 2023. The WHO fungal priority pathogens list as a game-changer. Nat Rev Microbiol 21:211–212.

8. Santiago-Tirado FH, Peng T, Yang M, Hang HC, Doering TL. 2015. A Single Protein S-acyl Transferase Acts through Diverse Substrates to Determine Cryptococcal Morphology, Stress Tolerance, and Pathogenic Outcome. PLoS Pathog 11:e1004908.

9. Nichols CB, Ost KS, Grogan DP, Pianalto K, Hasan S, Alspaugh JA. 2015. Impact of Protein Palmitoylation on the Virulence Potential of Cryptococcus neoformans. Eukaryot Cell 14:626–35.

10. Dettmann A, Heilig Y, Valerius O, Ludwig S, Seiler S. 2014. Fungal communication requires the MAK-2 pathway elements STE-20 and RAS-2, the NRC-1 adapter STE-50 and the MAP kinase scaffold HAM-5. PLoS Genet 10:e1004762.

11. Fischer MS, Glass NL. 2019. Communicate and Fuse: How Filamentous Fungi Establish and Maintain an Interconnected Mycelial Network. Front Microbiol 10:619.

12. Fu C, Sun S, Billmyre RB, Roach KC, Heitman J. 2015. Unisexual versus bisexual mating in Cryptococcus neoformans: Consequences and biological impacts. Fungal Genet Biol 78:65–75.

13. Sun S, Coelho MA, David-Palma M, Priest SJ, Heitman J. 2019. The Evolution of Sexual Reproduction and the Mating-Type Locus: Links to Pathogenesis of Cryptococcus Human Pathogenic Fungi. Annu Rev Genet 53:417–444.

14. Lin J, Idnurm A, Lin X. 2015. Morphology and its underlying genetic regulation impact the interaction between Cryptococcus neoformans and its hosts. Med Mycol 53:493–504.

15. Wang L, Zhai B, Lin X. 2012. The link between morphotype transition and virulence in Cryptococcus neoformans. PLoS Pathog 8:e1002765.

16. Zhai B, Wozniak KL, Masso-Silva J, Upadhyay S, Hole C, Rivera A, Wormley FL, Jr., Lin X. 2015. Development of protective inflammation and cell-mediated immunity against Cryptococcus neoformans after exposure to hyphal mutants. mBio 6:e01433–15.

17. Ning W, Jiang P, Guo Y, Wang C, Tan X, Zhang W, Peng D, Xue Y. 2021. GPS-Palm: a deep learning-based graphic presentation system for the prediction of S-palmitoylation sites in proteins. Brief Bioinform 22:1836–1847.

18. Fu C, Thielhelm TP, Heitman J. 2019. Unisexual reproduction promotes competition for mating partners in the global human fungal pathogen Cryptococcus deneoformans. PLoS Genet 15:e1008394.

19. Viviani MA, Esposto MC, Cogliati M, Montagna MT, Wickes BL. 2001. Isolation of a Cryptococcus neoformans serotype A MATa strain from the Italian environment. Med Mycol 39:383–6.

20. Son YE, Fu C, Jung WH, Oh SH, Kwak JH, Cardenas ME, Heitman J, Park HS. 2019. Pbp1-Interacting Protein Mkt1 Regulates Virulence and Sexual Reproduction in Cryptococcus neoformans. Front Cell Infect Microbiol 9:355.

21. Davidson RC, Nichols CB, Cox GM, Perfect JR, Heitman J. 2003. A MAP kinase cascade composed of cell type specific and non-specific elements controls mating and differentiation of the fungal pathogen Cryptococcus neoformans. Mol Microbiol 49:469–85.

22. Liu OW, Kelly MJ, Chow ED, Madhani HD. 2007. Parallel beta-helix proteins required for accurate capsule polysaccharide synthesis and virulence in the yeast Cryptococcus neoformans. Eukaryot Cell 6:630–40.

23. Kumar P, Heiss C, Santiago-Tirado FH, Black I, Azadi P, Doering TL. 2014. Pbx proteins in *Cryptococcus neoformans* cell wall remodeling and capsule assembly. Eukaryot Cell 13:560–71.

24. Denham ST, Verma S, Reynolds RC, Worne CL, Daugherty JM, Lane TE, Brown JCS. 2018. Regulated Release of Cryptococcal Polysaccharide Drives Virulence and Suppresses Immune Cell Infiltration into the Central Nervous System. Infect Immun 86.

25. Yoneda A, Doering TL. 2008. Regulation of *Cryptococcus neoformans* capsule size is mediated at the polymer level. Eukaryot Cell 7:546–9.

26. Fuchs BB, O’Brien E, Khoury JB, Mylonakis E. 2010. Methods for using Galleria mellonella as a model host to study fungal pathogenesis. Virulence 1:475–82.

27. Sun S, Priest SJ, Heitman J. 2019. Cryptococcus neoformans Mating and Genetic Crosses. Curr Protoc Microbiol 53:e75.

28. Nielsen K, Cox GM, Litvintseva AP, Mylonakis E, Malliaris SD, Benjamin DK, Jr., Giles SS, Mitchell TG, Casadevall A, Perfect JR, Heitman J. 2005. Cryptococcus neoformans alpha strains preferentially disseminate to the central nervous system during coinfection. Infect Immun 73:4922–33.

29. Lin X, Nielsen K, Patel S, Heitman J. 2008. Impact of mating type, serotype, and ploidy on the virulence of Cryptococcus neoformans. Infect Immun 76:2923–38.

30. Tian X, He GJ, Hu P, Chen L, Tao C, Cui YL, Shen L, Ke W, Xu H, Zhao Y, Xu Q, Bai F, Wu B, Yang E, Lin X, Wang L. 2018. Cryptococcus neoformans sexual reproduction is controlled by a quorum sensing peptide. Nat Microbiol 3:698–707.

31. Severin FF, Hyman AA. 2002. Pheromone induces programmed cell death in S. cerevisiae. Curr Biol 12:R233–5.

32. Wickes BL. 2002. The role of mating type and morphology in *Cryptococcus neoformans* pathogenesis. Int J Med Microbiol 292:313–29.

33. McClelland CM, Chang YC, Varma A, Kwon-Chung KJ. 2004. Uniqueness of the mating system in Cryptococcus neoformans. Trends Microbiol 12:208–12.

34. Kwon-Chung KJ, Fraser JA, Doering TL, Wang Z, Janbon G, Idnurm A, Bahn YS. 2014. *Cryptococcus neoformans* and *Cryptococcus gattii*, the etiologic agents of cryptococcosis. Cold Spring Harb Perspect Med 4:a019760.

35. Fu C, Heitman J. 2017. PRM1 and KAR5 function in cell-cell fusion and karyogamy to drive distinct bisexual and unisexual cycles in the Cryptococcus pathogenic species complex. PLoS Genet 13:e1007113.

36. Lee H, Chang YC, Kwon-Chung KJ. 2005. TUP1 disruption reveals biological differences between MATa and MATalpha strains of Cryptococcus neoformans. Mol Microbiol 55:1222–32.

37. Cruz MC, Fox DS, Heitman J. 2001. Calcineurin is required for hyphal elongation during mating and haploid fruiting in. Embo Journal 20:1020–1032.

38. Chow EW, Clancey SA, Billmyre RB, Averette AF, Granek JA, Mieczkowski P, Cardenas ME, Heitman J. 2017. Elucidation of the calcineurin-Crz1 stress response transcriptional network in the human fungal pathogen *Cryptococcus neoformans*. PLoS Genet 13:e1006667.

39. Chen Y, Toffaletti DL, Tenor JL, Litvintseva AP, Fang C, Mitchell TG, McDonald TR, Nielsen K, Boulware DR, Bicanic T, Perfect JR. 2014. The Cryptococcus neoformans transcriptome at the site of human meningitis. MBio 5:e01087–13.

40. Yu CH, Sephton-Clark P, Tenor JL, Toffaletti DL, Giamberardino C, Haverkamp M, Cuomo CA, Perfect JR. 2021. Gene Expression of Diverse Cryptococcus Isolates during Infection of the Human Central Nervous System. mBio 12:e0231321.

41. Camacho E, Casadevall A. 2018. Cryptococcal Traits Mediating Adherence to Biotic and Abiotic Surfaces. J Fungi (Basel) 4.

42. Huang MY, Joshi MB, Boucher MJ, Lee S, Loza LC, Gaylord EA, Doering TL, Madhani HD. 2022. Short homology-directed repair using optimized Cas9 in the pathogen Cryptococcus neoformans enables rapid gene deletion and tagging. Genetics 220.

43. Yount JS, Zhang MM, Hang HC. 2011. Visualization and Identification of Fatty Acylated Proteins Using Chemical Reporters. Curr Protoc Chem Biol 3:65–79.

44. Livak KJ, Schmittgen TD. 2001. Analysis of relative gene expression data using real-time quantitative PCR and the 2(-Delta Delta C(T)) Method. Methods 25:402–8.

45. Schindelin J, Arganda-Carreras I, Frise E, Kaynig V, Longair M, Pietzsch T, Preibisch S, Rueden C, Saalfeld S, Schmid B, Tinevez JY, White DJ, Hartenstein V, Eliceiri K, Tomancak P, Cardona A. 2012. Fiji: an open-source platform for biological-image analysis. Nat Methods 9:676-82.

46. Pierce CG, Uppuluri P, Tristan AR, Wormley FL, Jr., Mowat E, Ramage G, Lopez-Ribot JL. 2008. A simple and reproducible 96-well plate-based method for the formation of fungal biofilms and its application to antifungal susceptibility testing. Nat Protoc 3:1494–500.

47. Stempinski PR, Smith DFQ, Casadevall A. 2022. Cryptococcus neoformans Virulence Assay Using a Galleria mellonella Larvae Model System. Bio Protoc 12.

48. Tamura K, Stecher G, Kumar S. 2021. MEGA11: Molecular Evolutionary Genetics Analysis Version 11. Mol Biol Evol 38:3022–3027.

