## Supplemental Table S1 for "Phenotypic characterization of *HAM1*, a novel mating regulator of the fungal pathogen *Cryptococcus neoformans*"

| **Strain name** | **Drug used for selection** | **Reference or source** | **Strain description** |
| --- | --- | --- | --- |
| TDY450 | N/A | Doering lab | KN99a |
| TDY451 | N/A | Doering lab | KN99α |
| *ham1*Δα-NAT  (1-1) | Nourseothricin | This study | *ham1*Δ mutant in the MATα background (generated by biolistics) |
| *ham1*Δa-NAT  (1-2) | Nourseothricin | This study | *ham1*Δ mutant in the MATa background (generated by biolistics) |
| *ham1*Δa-NAT  (2-1) | Nourseothricin | This study | *ham1*Δ mutant in the MATa background (generated by crossing strain 1-1 with KN99a and picking spores) |
| C07 | Nourseothricin | Madhani lab deletion collection | *ham1*Δ mutant in the MATα background (from commercial deletion collection; generated by biolistics) |
| TDY1596 | G418 | Doering lab  Upadhya, R *et al*. Fungal genetics and biology (2017) | KN99α expressing mCherry from SH2 locus |
| TDY2096 | G418 | Doering lab | Same as TDY1596 but in MATa (obtained by crossing TDY1596 with KN99a and picking spores) |
| KN99a *CHS3*-mCherry (1-7) | Nourseothricin | Santiago-Tirado, FH *et al*. PLoS Pathogens (2015) | KN99a expressing *CHS3*-mCherry from its native locus |
| *ham1*Δα-NEO  (3-1) | G418 | This study | *ham1*Δ mutant in the KN99α background (generated by CRISPR) |
